# The N-terminal region of malaria vaccine candidate *Plasmodium falciparum* asparagine-rich merozoite antigen is immunodominant and targeted by polyreactive antibodies

**DOI:** 10.64898/2025.12.11.693633

**Authors:** Rolando Garza, Jeffrey M. Marchioni, Jared D. Honeycutt, Nicholas K. Hurlburt, Caroline Torres, Anakaren Garcia, Eva Loranc, Emily Yemington, Dalton Towers, Isaac Ssewanyana, Marie Pancera, Jason J. Lavinder, Prasanna Jagannathan, Bryan Greenhouse, Sebastiaan Bol, Evelien M. Bunnik

## Abstract

The development of malaria blood-stage vaccines has been hampered by sequence variation in many *Plasmodium falciparum* proteins involved in erythrocyte invasion. In the past few years, asparagine-rich merozoite antigen (PfARMA) has emerged as a potential vaccine candidate due to its low amino acid sequence diversity and the association between anti-PfARMA antibody levels and protection to malaria. Here, we used samples from *P. falciparum*-exposed individuals to study naturally acquired B cell and antibody responses to PfARMA. B cell responses to PfARMA were dominated by IgM^+^ B cells that recognized the N-terminal intrinsically disordered region 1 (IDR1) of PfARMA. A human monoclonal antibody (hmAb) to IDR1 was non-neutralizing, while a second hmAb binding to the folded domain showed weak neutralizing activity. Both PfARMA-specific plasma IgM and IgG responses predominately targeted IDR1 and their levels increased with *P. falciparum* exposure. However, in contrast to previous reports, these antibody responses did not correlate with protection in age and exposure-matched children. Interestingly, approximately 30% of unexposed individuals had IgG that also targeted IDR1 and was polyreactive, binding to regions with high asparagine content. Finally, we determined that PfARMA is located in or near PfEBA-175^+^ micronemes. These data suggest that while IgG to the folded domain of PfARMA may inhibit parasite growth, antibody responses to PfARMA are primarily directed to IDR1 and may not directly contribute to protection against malaria.

## INTRODUCTION

*Plasmodium falciparum* is the causative agent of more than 95% of malaria cases worldwide. In 2023, *P. falciparum* infections resulted in approximately 250 million cases of disease and 600,000 deaths, the majority of which were children under the age of five living in sub-Saharan Africa [1]. Despite progress in preventing malaria morbidity and mortality, new interventions are needed to achieve malaria eradication. A vaccine that protects against infection, disease, or transmission would be an important tool in this fight against malaria. The World Health Organization (WHO) has recommended two vaccines for the prevention of *P. falciparum* malaria in children living in malaria-endemic regions with moderate to high parasite transmission. Both vaccines, RTS,S/AS01 and R21/Matrix-M, target the sporozoite stage of *P. falciparum*, the form of the parasite transmitted from mosquitoes to humans. R21/Matrix-M is the first vaccine to meet the WHO’s goal of 75% vaccine efficacy over the first 12 months of follow up [2]. While this is a remarkable achievement, additional strategies will be needed to improve the efficacy and durability of malaria vaccines [3]. Adding a vaccine component that targets the *P. falciparum* blood-stage, which causes malaria pathogenesis, could help achieve this goal.

During the blood-stage, *P. falciparum* parasites invade erythrocytes and subsequently undergo asexual replication to produce 10–30 daughter parasites that will egress from the host cell and infect a new erythrocyte [4]. The erythrocyte invasion process is an important target of protective antibody responses [5–12]. As a result of intense immune pressure, many *P. falciparum* antigens that play a role during invasion have acquired genetic polymorphisms [13,14], hampering the development of vaccines that protect against globally circulating parasite strains. A notable exception is the highly conserved reticulocyte-binding protein homologue 5 (PfRh5), the leading candidate for a blood-stage vaccine [15–18]. Seroprevalence of PfRh5 antibody responses in *P. falciparum*-exposed individuals is low [19], possibly because the protein is essential for erythrocyte invasion and therefore carefully shielded from immune exposure. Another *P. falciparum* antigen that is highly conserved and has relatively low seroprevalence is asparagine-rich merozoite antigen (PfARMA; Pf3D7_1136200) [20,21]. These favorable characteristics suggest that PfARMA may be an additional candidate antigen for a blood-stage malaria vaccine.

PfARMA was identified as a potential novel vaccine candidate by Faith Osier *et al.*, who studied the correlation between antibody responses against combinations of five *P. falciparum* antigens and protection to malaria [20]. PfARMA was overrepresented in combinations of antigens that were predictive of high levels of protection. Subsequent studies have reported that plasma anti-PfARMA antibody reactivity correlates with protection to *P. falciparum* infection, malaria, or severe disease, depending on the study setup [21–23]. Moreover, Karamoko Niaré *et al*. showed that antisera generated via immunization of rabbits with full-length PfARMA protein induced Fc-mediated NK cell degranulation and inhibited *P. falciparum* growth *in vitro* [21]. However, it is unknown which region(s) of PfARMA are targeted by these inhibitory antibodies. Such information would inform the design of immunogens that focus the antibody response on the most relevant epitopes.

Here, we analyzed B cell and plasma antibody responses to PfARMA in *P. falciparum*-exposed individuals to determine the immunogenicity of distinct PfARMA fragments. Additionally, we isolated monoclonal antibodies to map the regions of PfARMA targeted by neutralizing antibodies. Finally, based on our observations in these experiments, we analyzed the binding characteristics of anti-PfARMA antibodies to better understand their polyreactive nature. Collectively, these analyses provide important new insights for the development of a PfARMA-based malaria vaccine.

## RESULTS

### The B cell response to PfARMA is dominated by IgM^+^ B cells that target an unstructured domain

To characterize the B cell response to PfARMA, we isolated PfARMA-specific B cells from five *P. falciparum*-exposed adults (**Table S1**) using a B cell tetramer constructed with full-length PfARMA protein. The PfARMA tetramer was mixed with tetramers for six other merozoite antigens: PfMSP1, PfMSP3, PfAMA1, Pf41, Pf113, and PfVFT. B cells with reactivity to any of the seven antigen tetramers were sorted into two populations based on isotype (IgM or IgG) (**Fig. S1**). Single B cells were stimulated in culture to differentiate into antibody-secreting cells, followed by screening of the culture supernatant for antigen specificity of the secreted human monoclonal antibodies (hmAbs). We identified a total of 32 hmAbs with confirmed PfARMA reactivity, with an overrepresentation of hmAbs derived from IgM^+^ B cells (n = 23, 72%) over IgG^+^ B cells (n = 9, 28%) (**Fig. 1A**). In contrast, the B cell responses to PfMSP3, PfAMA1, and Pf113 were relatively balanced (59 – 63% IgG^+^ B cells), while 98% of all B cells recognizing PfMSP1 were IgG^+^ (**Fig. 1A**). B cells targeting Pf41 and PfVFT were not included in this analysis because fewer than 10 hmAbs per antigen were isolated and isotype distributions were therefore not considered reliable (**Table S2**). Together, these results suggest that, unlike the response to other merozoite antigens, the humoral immune response to PfARMA is dominated by IgM^+^ B cells.

**Figure 1.**
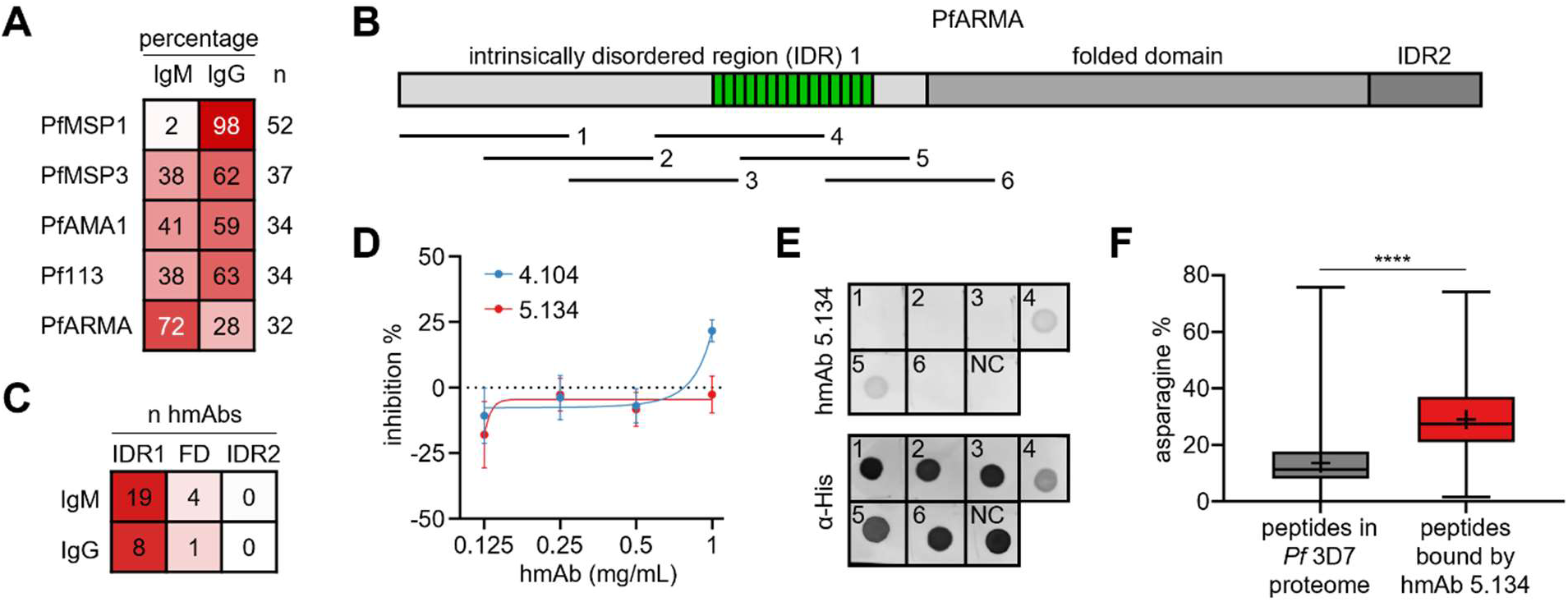
Isolation of human monoclonal antibodies to PfARMA. **A)** The percentage of IgM^+^ and IgG^+^ B cells with confirmed antigen-specificity obtained for PfARMA and four other merozoite proteins. **B)** Schematic of the predicted structural features of PfARMA. The asparagine-rich repeat region is shown in green. The six 100-amino acid (aa) long peptides with 50 aa overlap covering intrinsically disordered region (IDR) 1 are indicated below. **C)** The number of anti-PfARMA hmAbs targeting each of the three indicated regions of the protein. **D)** Percentage growth inhibition of *P. falciparum* strain 3D7 in *in vitro* cultures in the presence of hmAbs 5.134 and 4.104. Data points represent the average of three biological replicates, with the error bars showing the standard deviation. **E)** Dot-blot analysis showing reactivity of hmAb 5.134 to six IDR1 peptides and a negative control (NC) protein (PfAMA1). As a positive control for the presence of peptide and control protein, a second dot blot was stained in parallel with an anti-His antibody (bottom). **F)** The percentage of asparagine residues in overlapping 62-aa peptides covering the full *P. falciparum* (*Pf*) strain 3D7 proteome in the PhIP-seq library (n = 129,411) and among peptides bound by hmAb 5.134 (n = 303). Center line, median; box limits, upper and lower quartiles; whiskers, min/max values; +, mean. The difference between groups was tested using an unpaired student’s t-test. **** P < 0.0001.

Next, we sought to determine the regions of PfARMA that were bound by the anti-PfARMA hmAbs. The structure of PfARMA has yet to be resolved. Based on predictions of PfARMA structure by AlphaFold2 [24] and the presence of disordered protein regions by IUPred2 [25,26], the protein can be divided into three parts: intrinsically disordered region 1 (IDR1; amino acids 19 – 330), a folded domain (amino acids 331 – 595), and intrinsically disordered region 2 (IDR2; amino acids 596 – 656) (**Fig. 1B**). We produced recombinant PfARMA fragments corresponding to these three regions (**Fig. S2**) and measured reactivity of anti-PfARMA hmAbs to each fragment. The large majority of both IgM and IgG hmAbs (83% and 89%, respectively) bound to IDR1, with the remaining hmAbs recognizing the folded domain, and none binding to IDR2 (**Fig. 1C, Table S3**). Based on these results, we conclude that the B cell response to PfARMA is unusual, in that most PfARMA-specific B cells express IgM, and that both IgM^+^ and IgG^+^ B cells primarily recognize the same unstructured region of PfARMA.

### Monoclonal antibody 4.104 against the folded domain of PfARMA has limited neutralizing activity

To determine correlations between the regions of PfARMA targeted by hmAbs and their neutralizing activity, we aimed to produce recombinant versions of the hmAbs isolated by B cell sorting. We were able to generate recombinant IgG_1_ of two hmAbs: hmAb 5.134 binding to IDR1 and hmAb 4.104 targeting the folded domain (**Table S3, Fig. S3**). Both hmAbs were derived from IgM^+^ B cells, but were expressed as IgG_1_ to facilitate downstream analyses. hmAb 4.104 had low levels of somatic hypermutation (1% nucleotide changes), while hmAb 5.134 was substantially diversified from germline (11% nucleotide changes) (**Table S3**). We confirmed binding of hmAbs 4.104 and 5.134 to PfARMA using biolayer interferometry (**Fig. S4**). Both hmAbs showed weak binding with a moderate on-rate and fast off-rate.

In a growth-inhibition assay, hmAb 4.104 (targeting the folded domain) consistently showed approximately 20% inhibition at 1 mg/mL against *P. falciparum* strain 3D7, while 5.134 (targeting IDR1) did not show neutralizing activity at this concentration (**Fig. 1D, Table S4**). These results, albeit limited by the low number of hmAbs tested, may suggest that PfARMA-specific antibody responses against the folded domain can confer some level of parasite neutralization.

### Monoclonal antibody 5.134 binds the asparagine-rich repeat region

Although hmAb 5.134 did not show neutralizing activity, we were interested in understanding its binding properties, given that it targeted an intrinsically disordered region of PfARMA. To fine-map the epitope of hmAb 5.134, we expressed six 100-amino acid-long peptides with 50-amino acid overlaps spanning IDR1 (**Fig. 1B**). Peptides four (amino acids 169 – 268) and five (amino acids 219 – 318) were bound by hmAb 5.134 (**Fig. 1E**), suggesting that the epitope for hmAb 5.134 is located in the region shared by these two peptides (amino acids 219 – 268). This region is highly conserved and asparagine-rich, consisting of two major repeats (NNMNN and NNVNN) and several minor repeats (**Fig. S5**). Notably, it was previously reported that not a single mutation was found in this repeat region among 1,333 *P. falciparum* samples [21]. However, analysis of PfARMA protein sequences for the 18 *P. falciparum* strains from diverse geographical regions available on PlasmoDB revealed that 50% of the strains carry a deletion of one NNVNN repeat, and that three strains show additional genetic variation (**Fig. S5**).

PfARMA is not the only asparagine-rich protein in *P. falciparum*. To determine whether hmAb 5.134 showed cross-reactivity with other parasite proteins, we analyzed its reactivity profile using phage immunoprecipitation sequencing (PhIP-seq). The phage library contained ∼130,000 62-amino acid peptides with 37-amino acid overlap, covering the entire *P. falciparum* proteome [27]. In line with our previous observations, hmAb 5.134 bound three peptides covering the asparagine-rich repeat region in PfARMA IDR1 (**Table S5**). Additionally, we observed binding to 303 unique peptides from 158 *P. falciparum* strain 3D7 proteins (**Table S5**). On average, the asparagine content of these peptides was 29%, significantly higher than the average asparagine content of the full *P. falciparum* strain 3D7 proteome (14%, **Fig. 1F, Table S5**). Many of the non-PfARMA peptides bound by hmAb 5.134 contained repeats, but these were generally not the same as the major NNMNN and NNVNN repeats in PfARMA (**Table S5**). These results support our observation that hmAb 5.134 recognizes the asparagine-rich region in PfARMA (with up to 74% asparagine content) and suggest that hmAb 5.134 binds to asparagine-rich repeat regions without a specific sequence preference.

### PfARMA localizes to peripheral micronemes

It has previously been reported that PfARMA is located on the merozoite surface [21]. This localization would leave the protein exposed to the immune system, potentially resulting in a strong antibody response against PfARMA. However, the seroprevalence of anti-PfARMA antibodies is relatively low [20], which could point to localization in a subcellular structure, similar to PfRh5 [19]. We therefore decided to revisit the question of where PfARMA is localized in the merozoite. Despite its cross-reactivity with asparagine-rich regions in other *P. falciparum*proteins, we used hmAb 5.134 for this experiment, since hmAb 4.104 did not stain *P. falciparum* parasites. Although binding of hmAb 5.134 to non-PfARMA proteins in the immunofluorescence assay could not be completely ruled out, none of the hmAb 5.134 PhIP-seq protein hits are known to be located on the merozoite surface or in rhoptries, micronemes, or dense granules, which is the most likely location for PfARMA (**Table S4**). Additionally, hmAb 5.134 staining of *P. falciparum* segmented schizonts resulted in a staining pattern similar to that previously reported [21], and this staining was blocked by the addition of free recombinant PfARMA during primary antibody incubation (**Fig. S6**), suggesting that the signal is derived from specific PfARMA staining.

To determine the subcellular localization of PfARMA, we co-labeled segmented schizonts with hmAb 5.134 against PfARMA and a primary antibody targeting one of the following merozoite proteins with distinct localizations: PfMSP1 (surface membrane), PfAMA1 (micronemes), PfEBA-175 (micronemes), PfRON4 (rhoptry neck), PfRAP1 (rhoptry bulb), or PfRESA (dense granules) (**Fig. 2A, Fig. S7-13**). Although PfAMA1 and PfEBA-175 both reside in micronemes, these proteins mark different subsets of micronemes, of which PfAMA1 micronemes are localized closer to the apical end of the merozoite [28–30]. To quantify colocalization between PfARMA and the other proteins, we first calculated Pearson’s correlation coefficients between the signal intensity in the two channels for five representative parasites in each combination (**Table S6**). As a positive control, we used parasites that were stained with anti-PfRAP1 and two secondary antibodies with different fluorochromes. The positive control displayed almost perfect colocalization (median Pearson’s r, 0.97). PfARMA showed the highest Pearson’s correlation with PfEBA-175 (median Pearson’s r, 0.85) (**Fig. 2B**, top). The correlations between PfARMA and the other four proteins ranged from 0.67 to 0.75 (**Fig. 2B**, top), of which correlations with PfAMA1, PfRON4, and PfRAP1 were statistically significantly lower than the positive control. We also calculated Manders’ correlation coefficients (MCC) to capture the degree of overlap in signal between two channels. In line with the high correlation in signal intensity, nearly 100% of the PfEBA-175 signal overlapped with that of PfARMA (median MCC, 0.98), again the highest of all proteins and statistically significantly higher than for PfMSP1 and PfRESA (**Fig. 2B**, middle). Not all the PfARMA signal overlapped with PfEBA-175 (median MCC, 0.75) (**Fig. 2B**, bottom), which may indicate that its localization is more diffuse than PfEBA-175. Alternatively, this could be the result of hmAb 5.134 binding to non-PfARMA proteins. Collectively, these data suggest that PfARMA may not be found on the merozoite plasma membrane as previously reported, but is instead located inside or in close proximity to PfEBA-175-containing micronemes.

**Figure 2.**
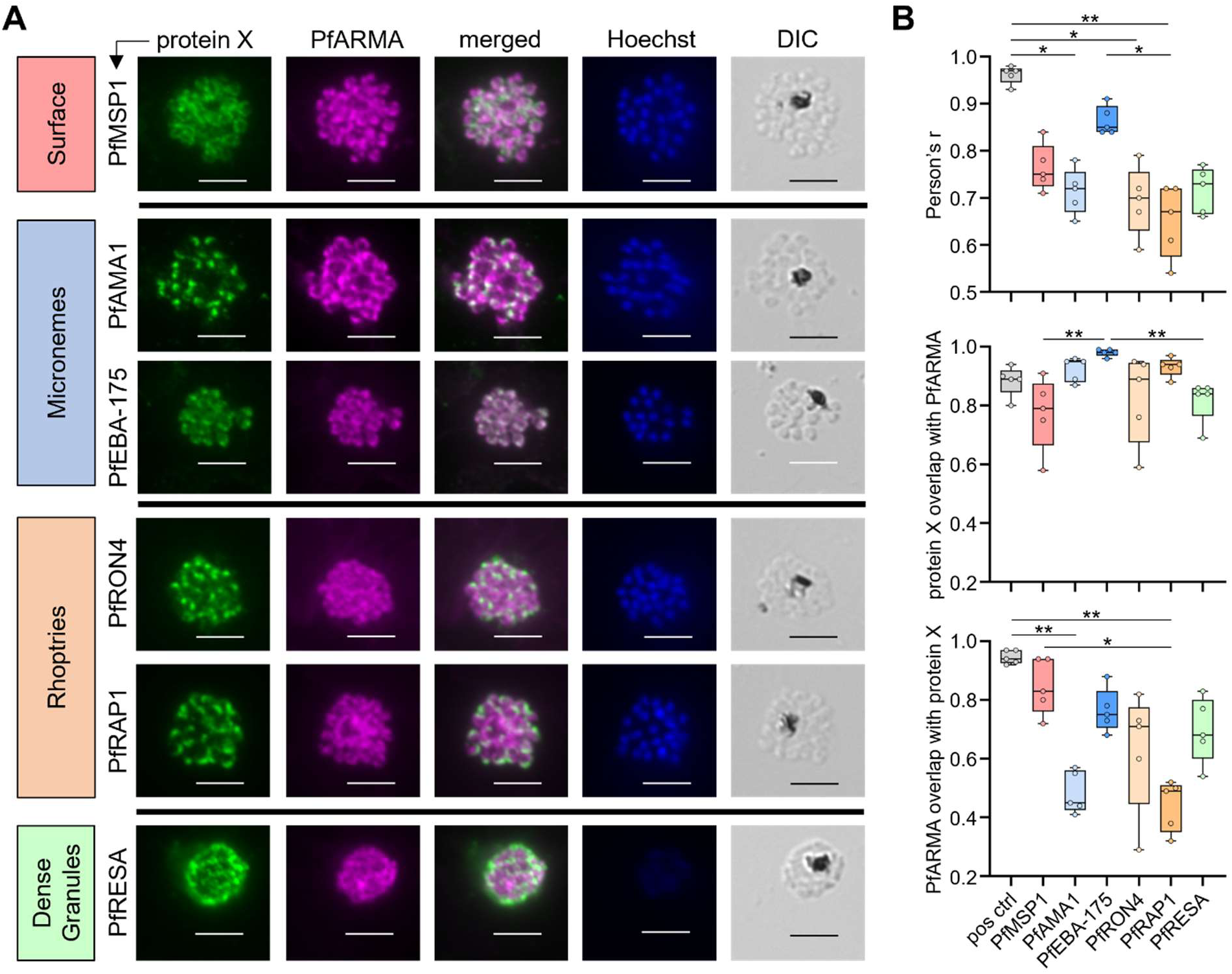
Colocalization of PfARMA with *P. falciparum* proteins present in distinct locations of the merozoite. **A)** Immunofluorescence images of segmented schizonts stained with Hoechst 33342 (DNA), hmAb 5.134 against PfARMA and an antibody against a second merozoite protein (protein X), indicated on the left. Scale bar denotes 5 µm. **B)** Colocalization analysis of PfARMA with merozoite protein X using five different parasites for each pairwise analysis. The positive control (pos ctrl) was performed using anti-PfRAP1 primary antibody and a mix of two secondary antibodies conjugated to different fluorophores. Top: Pearson’s correlation, middle: Manders’ coefficient 1 (fraction of second protein signal that overlaps with PfARMA signal), bottom: Manders’ coefficient 2 (fraction of PfARMA signal that overlaps with the second protein signal). Differences in correlation coefficients tested for statistical significance using a Kruskal-Wallis test, followed by comparisons between all pairs of groups using Dunn’s post-hoc test, which reports P values that have been corrected for multiple comparisons (n = 21). * P < 0.05; ** P < 0.01; *** P < 0.001.

### Plasma IgM and IgG levels to PfARMA increase with *P. falciparum* exposure but do not correlate with malaria susceptibility

Antibody responses to PfARMA have previously been associated with protection against *P. falciparum* malaria [20–23]. To confirm and extend these findings in an independent cohort, we studied the levels of anti-ARMA plasma IgM and IgG in *P. falciparum*-exposed individuals living in an area of high parasite transmission in Uganda. Since antibody levels against *P. falciparum* antigens typically increase with cumulative exposure, we selected two groups of children, who were matched for age (average, 6.2 and 6.1 years old, respectively; **Tables S1 and S7**) and exposure history. Based on the relative probability of experiencing symptomatic malaria when parasitemic with *P. falciparum*, these groups were defined as having low (n = 46) or moderate (n = 40) immunity against *P. falciparum* malaria. We also included adults, who have high levels of immunity (n = 54), to study whether anti-PfARMA antibody responses continue to develop with increased age and *P. falciparum* exposure. As negative controls, we used plasma samples from *P. falciparum*-naïve U.S. donors (n = 15).

As a positive control for *P. falciparum* exposure, we first measured plasma IgM and IgG reactivity to *P. falciparum* merozoite surface protein 1 (PfMSP1). As expected, the U.S. donors did not show plasma antibody reactivity to PfMSP1, confirming that they unlikely had significant prior exposure to *P. falciparum* (**Fig. 3A, Table S8**). For the subsequent analyses, we used the signal in plasma from these *P. falciparum*-naïve individuals to calculate the cutoff for seropositivity (three standard deviations above the average MFI). Among the *P. falciparum*-exposed individuals, 93% and 97% had plasma IgM and IgG reactivity to PfMSP1, respectively, with ≥90% seropositivity in every group (**Table S8**). For full-length PfARMA, we observed that 44%, 41%, and 68% of individuals with low, moderate, and high immunity to malaria had plasma IgM reactivity, respectively. For anti-PfARMA IgG reactivity, these numbers were 78%, 71%, and 88%, respectively. It should be noted that a subset of the *P. falciparum-*naïve individuals also had plasma IgG reactivity to full-length PfARMA (see below) and we therefore used a Gaussian mixed model to identify the subset of *P. falciparum-*naïve individuals without anti-PfARMA IgG reactivity to calculate the cutoff for seropositivity. PfARMA seropositivity in this study is higher than what has been previously reported, possibly as a result of the high sensitivity of the Luminex assay used for this experiment or a lower cutoff used. Importantly, there was no difference in anti-PfARMA IgM or IgG levels between the two groups of children with low and moderate immunity, while anti-PfARMA antibody levels in *P. falciparum-*exposed adults were significantly higher (**Fig. 3A, Table S8**). Based on these results, we conclude that although anti-PfARMA plasma IgM and IgG levels increase with cumulative exposure, they did not show an association with modeled immunity in age- and exposure-matched children in this cross-sectional analysis.

**Figure 3:**
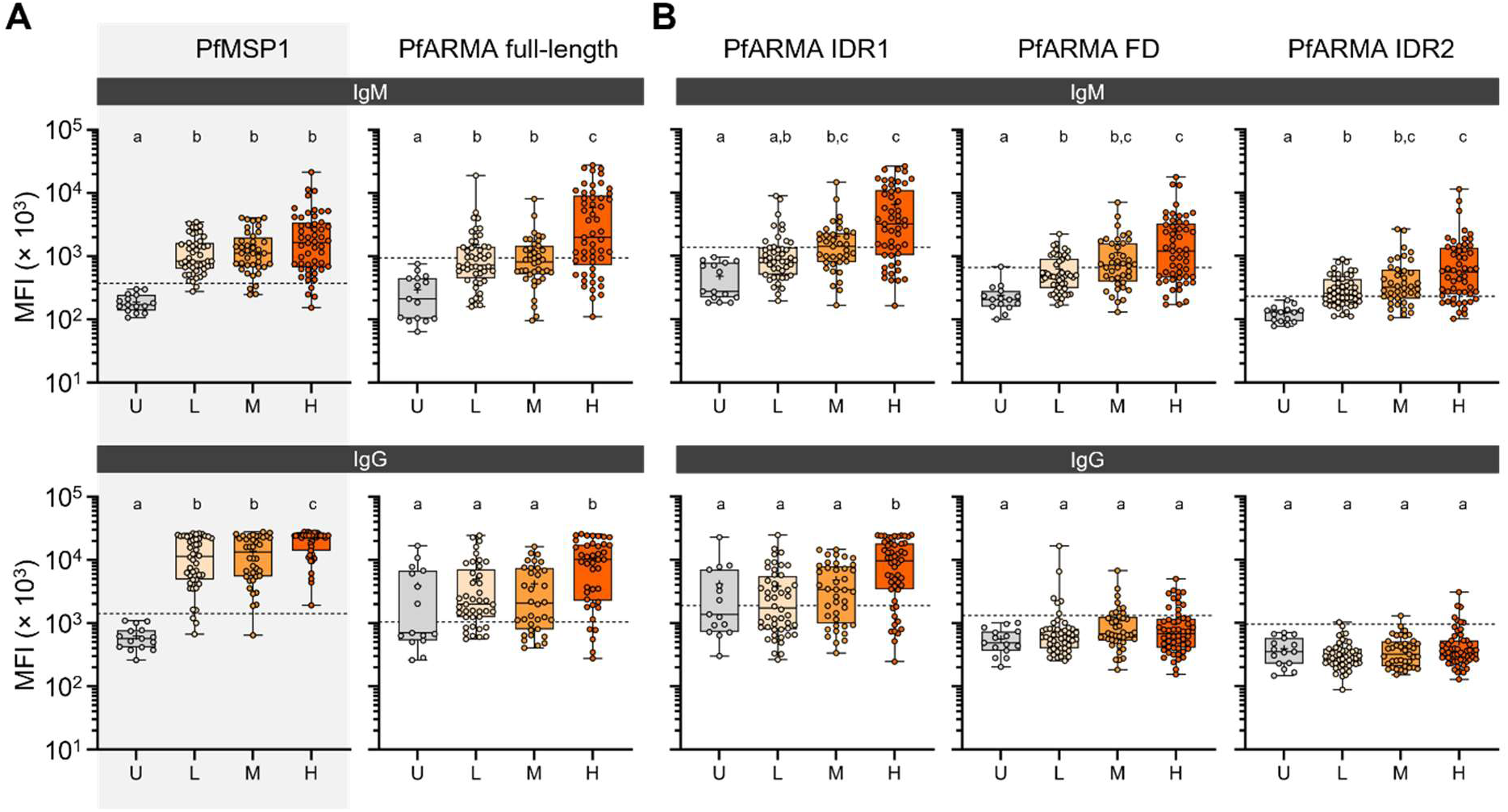
Plasma IgM and IgG reactivity to PfMSP1 and PfARMA. Plasma samples from *P. falciparum*-unexposed U.S. donors (U, n = 15) and *P. falciparum*-exposed individuals with low (L, n = 46), moderate (M, n = 40), and high (H, n = 54) immunity to malaria were measured in duplicate, with the average of both readings shown for (**A**) PfMSP1 and full-length PfARMA and (**B**) fragments of PfARMA. PfMSP1 was included as a control for *P. falciparum* exposure and is therefore shown on a gray background. The groups with low and medium immunity were matched for age and *P. falciparum* exposure history. The dashed horizontal line indicates the cutoff for seropositivity (three standard deviations above the average for the entire *P. falciparum*-naïve group, or for the subset of *P. falciparum*-naïve individuals without IgG reactivity for full-length PfARMA and IDR1). Center line, median; box limits, upper and lower quartiles; whiskers, min/max values; +, mean. Differences between groups were tested using a Kruskal-Wallis test, followed by comparisons between all pairs of groups using Dunn’s post-hoc test, which reports P values that have been corrected for multiple comparisons (n = 6). Within each graph, groups sharing the same letter are not statistically significantly different from each other, while groups with different letters are statistically significantly different (P < 0.05). No statistically significant differences were observed between the groups with low and medium immunity. IDR, intrinsically disordered region; FD, folded domain.

### Plasma anti-ARMA IgM and IgG are mainly directed against the first intrinsically disordered region

Since we observed that most B cells were reactive to the N-terminal IDR1 fragment of PfARMA (**Fig. 1C**), we were interested in determining whether plasma antibodies from *P. falciparum*-exposed individuals had the same reactivity profile. In line with our observations for B cells, plasma IgM and IgG levels were highest to intrinsically disordered region 1 (**Fig. 3B, Table S8**). Plasma IgM levels to IDR1, the folded domain, and IDR2 were higher in all three groups of *P. falciparum*-exposed individuals than in the unexposed controls, and a substantial proportion of the *P. falciparum*-exposed individuals (24% – 78%) was seropositive for IgM to these three regions (**Table S8**). In contrast, only a few individuals were seropositive for plasma IgG to the folded domain and IDR2, and no differences in IgG levels were observed between the *P. falciparum*-exposed and *P. falciparum*-naïve groups for these two fragments. Although more children with moderate immunity to malaria were seropositive for IgM to IDR1 and FD (51% and 55%, respectively) than among children with low immunity (24% and 31%, respectively), there was no difference in the median IgM or IgG reactivity to the three PfARMA fragments between the two groups of children. These data suggest that IDR1 is the immunodominant region of PfARMA, although plasma IgM to the folded domain and IDR2 were also detectable. IgM or IgG responses against none of these regions were associated with protection against malaria.

### *P. falciparum*-unexposed individuals show plasma IgG reactivity to PfARMA

Unexpectedly, we observed IgG reactivity to full-length PfARMA and IDR1 in six out of fifteen *P. falciparum*-naïve U.S. donors, at levels comparable to those in *P. falciparum*-exposed individuals (**Fig. 3A**). Among these were four healthy blood donors (out of eight, 50%) and two convalescent COVID-19 patients sampled within a month after symptom onset (out of seven, 29%). Their plasma IgG showed reactivity to PfARMA, but not to six other merozoite antigens (**Fig. 4A, Table S9**). Although quite unlikely, a possible explanation for this observation could be that these individuals had an autoimmune disease, since prior studies have reported that patients with autoimmune disease can develop antibodies that bind to *P. falciparum* antigens [31–33]. To determine if anti-PfARMA IgG reactivity is a prominent feature of the autoantibody responses in patients with antibody-mediated autoimmune disease, we measured antibody reactivity to the seven *P. falciparum* proteins in plasma from ten systemic lupus erythematosus (SLE) patients and eight seropositive rheumatoid arthritis (RA) patients with active disease. Two SLE patients (20%) and two RA patients (25%) showed plasma anti-PfARMA IgG reactivity (MFI > 2.6 × 10^3^). Interestingly, one SLE patient showed IgG reactivity to five other merozoite proteins, but not PfARMA (**Table S9**). These results indicate that a substantial fraction (∼30%) of *P. falciparum*-naïve individuals show IgG reactivity to PfARMA, and suggest this is not related to recent infection or an underlying autoimmune disorder.

**Figure 4:**
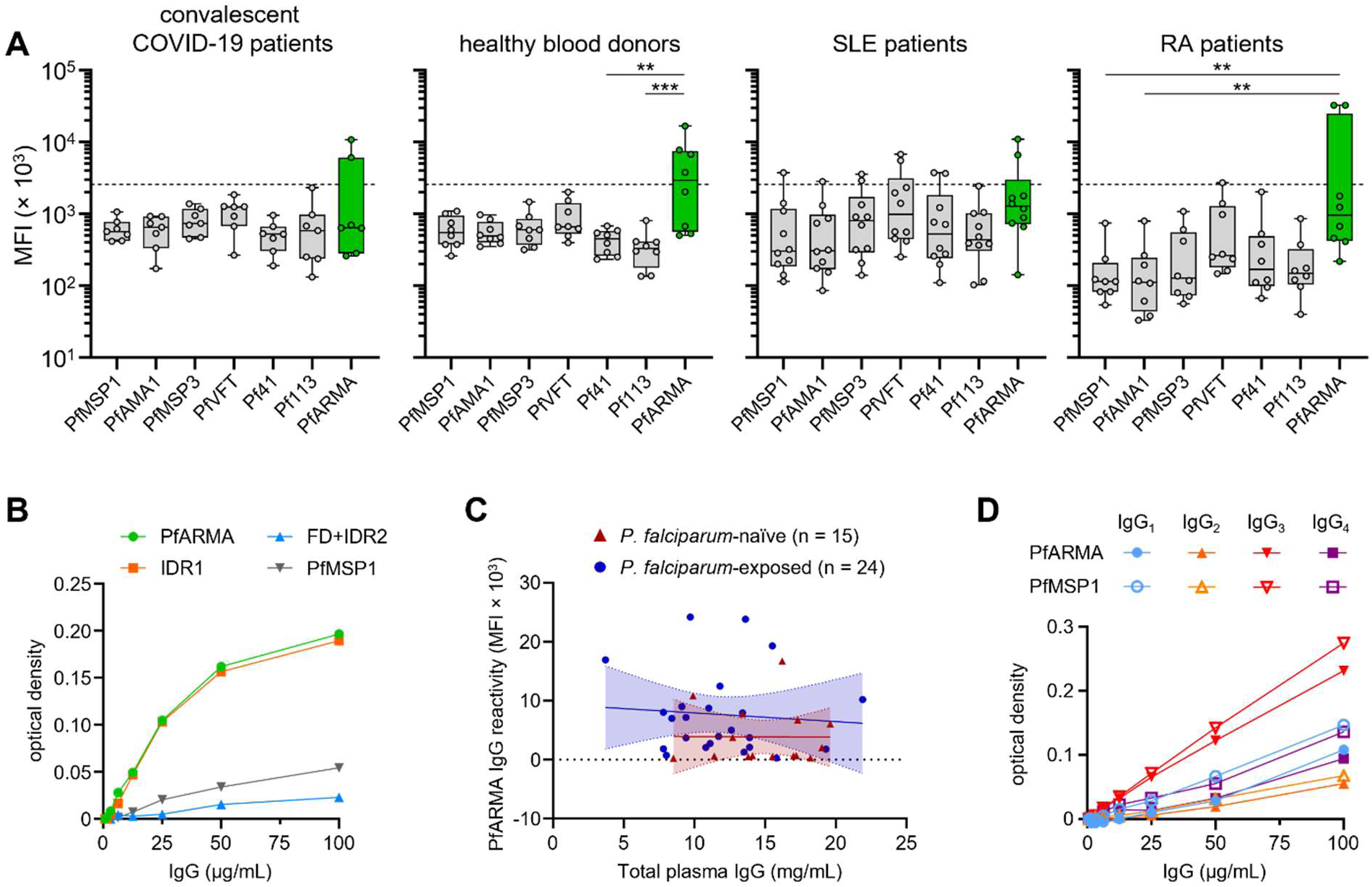
Plasma IgG reactivity to PfARMA and six other merozoite antigens in *P. falciparum*-naïve individuals. **A)** Plasma samples were obtained from *P. falciparum*-unexposed individuals living in the U.S. who were convalescent COVID-19 patients (n = 7), healthy blood donors (n = 8), systemic lupus erythematosus (SLE) patients (n = 10), or rheumatoid arthritis (RA) patients (n = 8). In all plots, the horizontal dashed line indicates the cut-off for reactivity to PfARMA (MFI = 2.6 × 10^3^), which equals the average MFI + three standard deviations of reactivity to the other six merozoite antigens among samples from non-autoimmune donors (convalescent COVID-19 patients and healthy blood donors). Differences in reactivity to the various merozoite antigens were tested for statistical significance using a Kruskal-Wallis test, followed by comparisons between PfARMA and all other antigens using Dunn’s post-hoc test, which reports P values that have been corrected for multiple comparisons (n = 6). ** P < 0.01; *** P < 0.001. **B)** Dose-dependent reactivity of IgG purified from plasma of a *P. falciparum*-naïve U.S. donor with high anti-PfARMA IgG reactivity to PfMSP1, full-length PfARMA, and fragments of PfARMA, as measured by ELISA. IDR, intrinsically disordered region; FD, folded domain. **C)** Total plasma IgG concentration plotted against PfARMA IgG reactivity for *P. falciparum*-naïve individuals and *P. falciparum*-exposed individuals. The line shows the best-fit using simple linear regression, with 95% confidence intervals. **D)** Reactivity of recombinant IgG1, IgG2, IgG3, and IgG4 with specificity for a different *P. falciparum* antigen to PfARMA and PfMSP1.

### The interaction of plasma IgG with PfARMA is not the result of non-specific binding through the Fc-tail

To study the specificity of the interaction between plasma IgG and PfARMA, we first purified total IgG from the plasma of a *P. falciparum*-naïve U.S. donor with high plasma IgG reactivity to PfARMA and measured binding to PfARMA in an ELISA. We observed dose-dependent binding of purified total plasma IgG to full-length PfARMA and IDR1, but not to the folded domain or IDR2 (**Fig. 4B, Table S10**). These results confirm that IgG directly interacts with PfARMA in the absence of other plasma components. However, the dose-dependent binding of IgG to PfARMA could be the result of non-specific interactions, for example mediated by the Fc tail of IgG. If this were the case, we would expect that higher total plasma IgG concentrations would result in higher PfARMA binding. To test this hypothesis, we quantified total plasma IgG in 15 *P. falciparum*-naïve U.S. adults and 24 *P. falciparum*-exposed adults. While the *P. falciparum*-naïve individuals had higher total plasma IgG concentrations than the *P. falciparum*-exposed individuals (**Fig. S14**), we did not observe a correlation between plasma IgG concentration and PfARMA reactivity in either group (**Fig. 4C, Table S11**). Finally, we tested whether PfARMA interacts non-specifically with a certain IgG isotype, which could have been missed in our experiments with total plasma IgG. We generated recombinant IgG_1_, IgG_2_, IgG_3_, and IgG_4_ versions of the same non-PfARMA-specific monoclonal antibody. Each of the four IgG isotypes showed the same level of reactivity to PfARMA and PfMSP1 (**Fig. 4D, Table S12**). Together, these results demonstrate that plasma IgG from *P. falciparum*-naïve individuals does not bind non-specifically to PfARMA via the heavy chain constant region, but rather specifically through the variable regions.

### Anti-PfARMA plasma IgG has polyreactive properties

Although we did not find that patients with antibody-mediated autoimmune disease were more likely to have anti-PfARMA plasma reactivity, autoantibodies are also common in healthy individuals [34–36]. We therefore hypothesized that anti-PfARMA IgG reactivity in *P. falciparum*-naïve individuals may arise through cross-reactivity of autoantibodies with PfARMA. To test this hypothesis, we purified autoantibodies from the three *P. falciparum*-naïve healthy blood donors with the highest anti-PfARMA plasma IgG reactivity. We first isolated total plasma IgG, followed by affinity-purification of autoantibodies using human cell line lysate (**Fig. 5A**). When tested at the same concentration, the autoantibody fractions on average showed a twofold increase in binding to PfARMA as compared to the flowthrough fractions. However, the autoantibody fractions consisted of only 1 – 2% of total plasma IgG, and most of the PfARMA reactivity thus remained in the flowthrough fraction (**Fig. 5B, Table S13**). Interestingly, in comparison to PfARMA, the autoantibody fractions showed a larger enrichment for reactivity to all other merozoite antigens, in particular PfAMA1 (47-fold increase in reactivity). In line with the reactivity profile of total plasma IgG, the anti-PfARMA autoantibody fractions bound to IDR1 (**Fig. 5B**). These results show that autoreactive IgG can cross-react with PfARMA, but this constitutes a very small fraction of anti-PfARMA plasma IgG.

**Figure 5:**
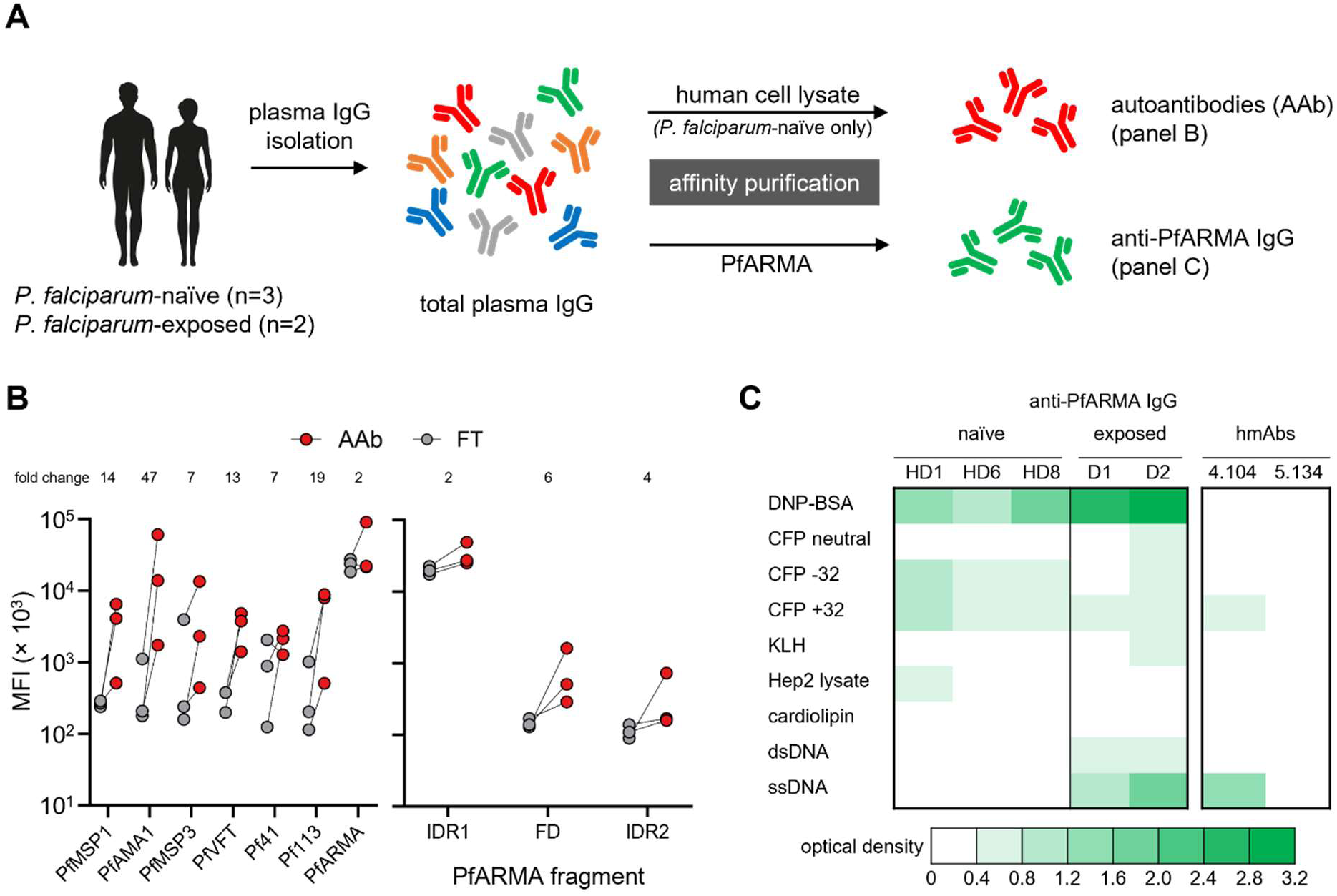
Polyreactivity of anti-PfARMA antibodies. **A)** Overview of the affinity purification of autoantibodies and anti-PfARMA IgG from plasma IgG from *P. falciparum*-naïve and *P. falciparum*-exposed individuals. **B)** Reactivity of affinity-purified autoantibodies (AAb) and the flowthrough (FT) fraction from plasma IgG of *P. falciparum*-naïve individuals, both tested at 20 µg/mL, to a panel of seven merozoite antigens (left) and PfARMA fragments (right). The average fold change in MFI in the autoantibody fractions relative to the flowthrough fractions is indicated in the top. **C)** Heatmap showing binding of anti-PfARMA plasma IgG isolated from *P. falciparum*-naïve and *P. falciparum*-exposed individuals by affinity purification, as well as anti-PfARMA hmAbs, to (macro)molecules with properties often recognized by auto- or polyreactive antibodies, measured by ELISA. Values are color-coded based on the optical density (OD) after subtraction of the background signal. OD = 0.4 was used as the cutoff for binding. DNP-BSA, dinitrophenol-bovine serum albumin; CFP, cyan-fluorescent protein; KLH, keyhole limpet hemocyanin; dsDNA, double-stranded DNA; ssDNA, single-stranded DNA.

Having observed that most anti-PfARMA plasma IgG does not cross-react with autoantigens, we wondered if anti-PfARMA IgG had certain binding properties in common that could explain their reactivity to PfARMA. To address this question, we affinity-purified anti-PfARMA IgG from three *P. falciparum*-naïve individuals with the highest PfARMA plasma IgG reactivity. Both the anti-PfARMA and flowthrough IgG fractions were tested for binding to a collection of proteins and other (macro)molecules with structural or chemical properties that are frequently bound by polyreactive or autoreactive antibodies [37–42]. We also included anti-PfARMA and flowthrough IgG fractions from two *P. falciparum*-exposed adults and hmAbs 4.104 and 5.134 to allow for a comparison of antibody binding characteristics between unexposed and exposed donors. Affinity-purified anti-PfARMA IgG fractions from all donors bound to dinitrophenol-conjugated bovine serum albumin (DNP-BSA), an indicator of antibody polyreactivity [43,44] (**Fig. 5C, Table S14**). Anti-PfARMA IgG from *P. falciparum*-naïve individuals showed binding to strongly positively charged (+32) and negatively charged (−32) cyan-fluorescent protein (CFP), but not to uncharged CFP, which was included as a control (**Fig. 5C, Table S14**). We also observed binding to Hep2 lysate in one of the samples from unexposed donors, indicative of reactivity to autoantigens, confirming that some anti-PfARMA IgG is cross-reactive with autoantigens. Conversely, anti-PfARMA IgG from *P. falciparum*-exposed individuals, but not from unexposed donors, was reactive with double-stranded and single-stranded DNA, as was hmAb 4.104, which could suggest that this signal comes from antibodies targeting the folded domain. hmAb 5.134 did not show a unique signal that can be associated with IgG targeting the asparagine repeat region. Collectively, these data suggest that anti-PfARMA IgG has polyreactive properties, in part through interactions with charged amino acids.

### Polyreactive antibodies bind to the asparagine-rich region in PfARMA

Having established that anti-PfARMA IgG from *P. falciparum*-naïve individuals predominantly targets IDR1 of PfARMA and interacts with charged amino acids (aa), we were interested in determining where in the 330 aa-long IDR1 these polyreactive antibodies bind. IDR1 contains a repeat region (aa 205 – 287) in which 76% of all amino acid residues are the polar amino acid residue asparagine (**Fig. S15**). In addition, it contains two strongly negatively charged regions with a net charge per residue (NCPR) smaller than −0.5: aa 83 – 102 (NCPR = −0.8) and aa 169 – 181 (NCPR = −0.53) (**Fig. 6A, Fig. S15**). To determine if these polar and charged regions in IDR1 were the main targets of PfARMA-reactive IgG, we used phage immunoprecipitation sequencing (PhIP-seq) to analyze plasma samples from five *P. falciparum*-naïve adults and 24 *P. falciparum*-exposed children and adults, who were selected based on high PfARMA IgG reactivity. We reasoned that the PhIP-seq platform would be well-suited for measuring plasma IgG reactivity to peptides from IDR1, as we had already confirmed that hmAb 5.134 bound to peptides derived from the asparagine-rich repeat region (**Table S4**). We observed reactivity with PfARMA peptides in two of the five *P. falciparum*-naïve individuals, both showing binding to two overlapping peptides from the asparagine-rich repeat region (**Fig. 6A, Table S15**). Additionally, we detected binding of one PfARMA peptide in plasma samples from four of 25 *P. falciparum*-exposed individuals. In one person, this was a peptide derived from the asparagine-rich repeat region, while the other three individuals showed enrichment for a peptide containing the shorter negatively charged patch (**Fig. 6A, Table S15**). In most samples, we did not detect IgG binding to PfARMA peptides, despite plasma IgG reactivity to PfARMA IDR1 (**Fig. 3B**). The overall peptide enrichment profiles in these samples were dominated by reads mapping to a small number of *P. falciparum* antigens. As a result, the sequencing depth may not have been sufficient to detect antibody reactivity to PfARMA. We therefore also analyzed reactivity to non-PfARMA peptides that were bound by hmAb 5.134 (**Table S4**), since antibodies that recognize these peptides may cross-react with PfARMA.

**Figure 6:**
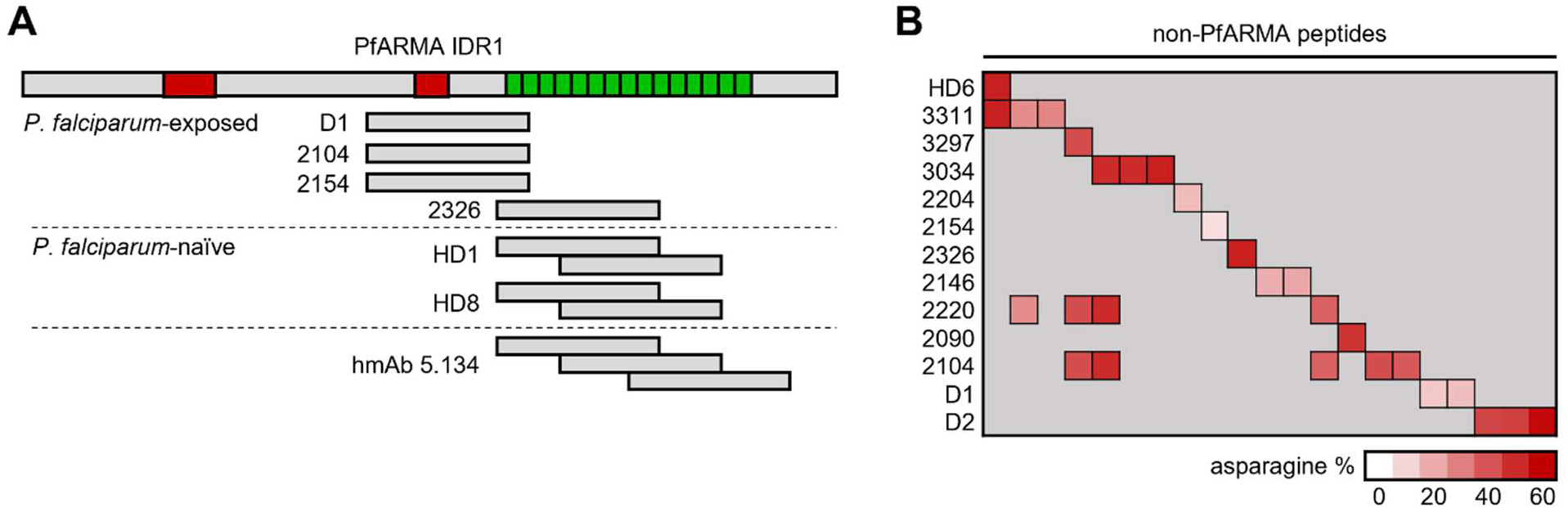
Reactivity to asparagine-rich peptides from PfARMA and other *P. falciparum* antigens. **A)** PfARMA peptides recognized by plasma IgG from *P. falciparum*-exposed and *P. falciparum*-naïve individuals. All peptides were located in intrinsically disordered region (IDR) 1 and overlapped either a negatively charged patch (shown in red) or the asparagine-rich repeat region (shown in green). **B)** Non-PfARMA-derived peptides that were bound by hmAb 5.134 and one or more plasma IgG samples. The red shading indicates the percentage of asparagine residues in the peptides. Absence of reactivity to peptides is indicated with gray shading.

In one additional *P. falciparum*-naïve individual and 12 *P. falciparum-*exposed donors (which includes the four individuals with detected reactivity to PfARMA peptides), significant reactivity was observed to 21 non-PfARMA peptides that had also been bound by hmAb 5.134 (**Fig. 6B, Table S15**). The average asparagine content of the 21 non-PfARMA peptides recognized by these samples was high (36%) in comparison to the average asparagine content of the full *P. falciparum* proteome (14%, **Fig. 1F**). Collectively, these results show that antibodies that interact with asparagine-rich regions were detected in approximately 50% of *P. falciparum*-naïve and *P. falciparum*-exposed individuals, and that such antibodies mediate PfARMA reactivity in plasma from *P. falciparum*-naïve individuals.

## DISCUSSION

The highly conserved *P. falciparum* protein PfARMA has emerged as a potential candidate for a malaria vaccine against the parasite’s blood-stage. Several studies have reported a correlation between the presence of anti-PfARMA plasma antibodies and protection from *P. falciparum* malaria [20–23]. Here, we aimed to map the antigenic landscape of PfARMA by measuring PfARMA-specific B cell and plasma antibody responses to different parts of the protein. We observed that the intrinsically disordered N-terminal domain is immunodominant in *P. falciparum*-exposed individuals living in a region of high parasite transmission. Unexpectedly, we also observed antibody reactivity to this region in *P. falciparum*-naïve individuals living in the United States. We determined that this reactivity is mediated by polyreactive antibodies that bind to a highly asparagine-rich region.

In contrast to six other merozoite antigens that were used for the isolation of antigen-specific B cells, the response to PfARMA in *P. falciparum*-exposed individuals was dominated by IgM^+^ B cells. This observation is reminiscent of the B cell response to PfMSP1 in young children, who have more IgM^+^ than IgG^+^ PfMSP1-specific B cells [45,46]. The samples that were used for the experiments in the current study were obtained from adults with life-long exposure to *P. falciparum*, in whom the PfMSP1-specific B cell response was almost entirely made up of IgG^+^ B cells, as reported previously [45,46]. One possible explanation for this difference in B cell receptor isotype between B cells recognizing PfARMA and PfMSP1 in adults is that a strong IgM^+^ B cell response is indicative of low cumulative levels of exposure. For PfMSP1, this would fit with an IgM^+^ B cell response in young children, followed by a shift towards IgG^+^ B cell responses later in life. For PfARMA, this would be in line with the relatively low immunogenicity of PfARMA, resulting in low PfARMA seroprevalence in *P. falciparum*-exposed individuals, even when living under high parasite transmission. An alternative explanation for detecting a more prominent IgM^+^ than IgG^+^ B cell response to PfARMA could be that many of these cells are low-affinity IgM^+^ B cells with some level of polyreactivity. Most PfARMA-specific B cells target intrinsically disordered region (IDR) 1, which contains the asparagine-rich repeat region bound by polyreactive antibodies. Indeed, mAb 5.134 was derived from an IgM^+^ B cell, showed weak binding in biolayer interferometry experiments, and is cross-reactive with asparagine-rich peptides in other *P. falciparum* proteins. However, the proportion of IgG^+^ B cells that bind to IDR1 is similar to that among IgM^+^ B cells, indicating that the distribution of B cells targeting the different regions does not change as the B cell response matures. At the plasma IgM and IgG level, PfARMA IDR1 was also the main target. It therefore seems likely that IDR1 is intrinsically immunodominant. In the context of vaccine development, a PfARMA variant that does not contain IDR1 could be immunogenic and elicit immune responses to the folded region or IDR2.

PfARMA IDR1 contains low-complexity sequences and an asparagine-rich repeat region. Protein repeats and low-complexity regions have been implicated in immune evasion by *P. falciparum* [47,48]. By measuring IgG responses to linear B cell epitopes of 37 *P. falciparum* proteins, Hou *et al.* determined that repetitive, low-complexity regions and glutamate-rich motifs are highly antigenic, but that IgGs targeting these regions do not inhibit invasion of erythrocytes by *P. falciparum* [48]. Similarly, vaccination with PfCSP containing the NANP/NVDP repeat region resulted in immune responses that predominantly target this repeat region, with limited immune responses to other domains of the protein [49]. In contrast, vaccination with PfCSP lacking the repeat region resulted in the expansion of immune responses to subdominant epitopes, which conferred better protection to malaria [49]. Mechanistically, the strong stimulation of B cells by repetitive antigens may elicit a rapid and dominant plasmablast response. These metabolically hyperactive plasmablasts can inhibit geminal center reactions and impair humoral immune responses to the parasite [50]. As such, it is possible that IDR1 of PfARMA functions to divert immune responses away from inhibitory epitopes and inhibit the formation of long-lived B cell responses against functional regions of PfARMA or other proteins. Interestingly, Hou *et al.* reported that highly antigenic, low-complexity epitopes had low levels of sequence variation between distinct *P. falciparum* strains [48], similar to the highly conserved asparagine-rich repeat region in PfARMA. The value of these epitopes to the parasite as an immune evasion strategy may explain their low propensity to genetic mutations despite their high immunogenicity.

In this study, we did not observe a correlation between PfARMA plasma IgM or IgG levels and protection in age and exposure-matched children. This is seemingly at odds with other reports suggesting a strong correlation between the presence of anti-PfARMA plasma IgM and IgG reactivity and protection to *P. falciparum* malaria [20–23]. A potential explanation could be that PfARMA plasma IgGs are short-lived and that higher PfARMA plasma IgG reactivity is indicative of more recent exposure. Yman *et al.* reported that one year after infection anti-PfARMA IgG levels were reduced by more than 90%, and that IgG responses against PfARMA could be used to determine with approximately 75% accuracy whether a person had experienced a *P. falciparum* infection in the preceding three months [51]. The fast waning of PfARMA antibody responses and our observation that antibodies against PfARMA are predominantly directed to the repeat-containing IDR1 region are in line with a report that IgG responses to repeat-containing *P. falciparum* antigens are short-lived and dependent on exposure [27]. In general, the levels and breadth of antibody responses are highest shortly after infection, and *P. falciparum* infection is no exception to this rule [52]. We therefore propose that individuals with higher plasma IgM or IgG reactivity to PfARMA may be protected from subsequent symptomatic infection due to broad, recently acquired antibody responses that target multiple parasite antigens, not because of the presence of anti-PfARMA antibodies by itself.

Our results show that anti-PfARMA IgGs have polyreactive properties, including reactivity to self-antigens. Many studies have reported autoantibody responses in *P. falciparum*-exposed individuals, and their presence is associated with both pathology and protection [31,53–58]. Additionally, plasma antibodies from patients with autoimmune diseases can bind *P. falciparum* antigens and autoantibodies from these patients possess anti-*P. falciparum* activity *in vitro* [32,33]. Hagadorn *et al.* [31] performed the most comprehensive analysis to date of autoreactive antibodies isolated from *P. falciparum*-exposed individuals, showing low autoantibody reactivity to PfARMA (bottom 50% of 668 *P. falciparum* antigens tested). Similarly, our results show some reactivity to PfARMA in autoantigen-affinity-purified IgG, and conversely, some reactivity to Hep2 lysate (an indicator of autoantibodies) in PfARMA-affinity-purified IgG samples. However, other binding properties of anti-PfARMA IgG were more striking, in particular the strong reactivity to dinitrophenol conjugated to bovine serum albumin (DNP-BSA). DNP is a synthetic molecule not found in the environment, and any antibody reactivity to DNP can therefore be considered to reflect polyreactive properties [43,44]. In our experience, DNP-binding antibodies are often correlated with poor biophysical properties, such as the tendency to aggregate or a low melting temperature. DNP reactivity in affinity-purified anti-PfARMA IgG samples can thus be indicative of both direct DNP binding and non-specific enrichment during the affinity purification process as a result of, for example, aggregation on the column. In addition to DNP, all samples showed binding to several other macromolecules, providing additional evidence for their polyreactive nature.

The reactivity profiles of anti-PfARMA IgG isolated from *P. falciparum*-naïve and *P. falciparum*-exposed individuals also showed differences, such as DNA binding in samples from *P. falciparum*-exposed but not *P. falciparum*-naïve individuals. These differences in reactivity profiles may be related to the specific epitopes that are recognized by anti-PfARMA antibodies. Our PhIP-seq results suggest that anti-PfARMA IgG from *P. falciparum*-naïve individuals may primarily target the asparagine-rich repeat region, while anti-PfARMA IgG from *P. falciparum*-exposed individuals also recognize an epitope directly upstream of this repeat region. Although mAb 5.134, targeting the asparagine-repeat region, did not show neutralizing activity in an *in vitro* growth inhibition assay, it remains to be determined whether antibodies to this upstream region can neutralize the parasite. Besides neutralization, antibody inhibition of *P. falciparum* merozoites can also occur through Fc-mediated effector functions, which to some extent depends on the localization of the antigen. Opsonic phagocytosis activity is greatest for antibodies against merozoite surface antigens, whereas complement-fixing activity is greater for antibodies against rhoptry and microneme antigens [9,12,59]. Our results show that PfARMA is most likely localized to PfEBA-175^+^ micronemes, suggesting that Fc-dependent mechanisms may be less effective *in vivo* due to the short time frame that PfARMA will be accessible on the parasite surface and the space constraints at the parasite-erythrocyte interface. However, it has been reported that anti-PfARMA IgG can exert parasite inhibition through Fc-mediated degranulation of natural killer cells [21]. Fc-mediated effector functions of anti-PfARMA IgG will thus need to be further explored.

## CONCLUSIONS

Our data suggest that the N-terminal IDR1 of PfARMA is immunodominant and is targeted by antibodies with polyreactive properties that bind to an asparagine-rich repeat region. Although our conclusions are limited by the small number of hmAbs tested, we did not observe neutralizing activity to this repeat region. We also did not observe a correlation between plasma anti-PfARMA IgM or IgG levels and protection from malaria. High plasma anti-PfARMA IgG levels, predominantly targeting IDR1, may primarily be an indicator of recent infection and as such be indirectly correlated with protection from infection or disease. On the other hand, a hmAb directed to the folded domain of PfARMA weakly inhibited parasite growth. Based on these immunological properties, our results suggest that vaccine development should focus on the folded domain of PfARMA. However, more research into epitope-specific responses and effector functions of inhibitory antibodies is needed to establish the value of PfARMA as a blood-stage malaria vaccine candidate.

## METHODS

### Ethics approval statement

*P. falciparum*-exposed individuals were enrolled in the Program for Resistance, Immunology, Surveillance, and Modeling of Malaria (PRISM) Cohort, and provided written consent for the use of their samples for research. These cohort studies were approved by the Makerere University School of Medicine Research and Ethics Committee (SOMREC) and the University of California, San Francisco Human Research Protection Program & Institutional Review Board. The use of cohort samples was approved by the Institutional Review Board of the University of Texas Health Science Center at San Antonio.

Blood donors consented to the use of their blood for research. The use of samples from anonymous blood donors was not considered human research by the Institutional Review Board of the University of Texas Health Science Center at San Antonio due to the lack of any identifiable information and was therefore exempt from review.

Convalescent COVID-19 patients, SLE patients, and RA patients provided written informed consent prior to specimen collection for the collection of associated clinical information and use of clinical specimens for research. These studies were reviewed and approved by the University of Texas Health Science Center at San Antonio Institutional Review Board.

### Sample selection

*P. falciparum*-exposed individuals originated from Tororo District in Eastern Uganda (n=66; annual entomological inoculation rate in the region estimated at 125 infectious bites per person per year) or Kanungu District in Western Uganda (n=78; annual entomological inoculation rate in the region estimated at 26.6 infectious bites per person per year) [60], and were selected for inclusion into this study based on age and relative probability of experiencing malaria when infected with *P. falciparum*. Specifically, generalized additive models with individual-level random effects were used to model the binary probability of being symptomatic given infection, adjusted by age (smooth), estimated exposure based on site (categorical), and log entomological inoculation rate (smooth). Individuals with the highest random effects (indicating a higher probability of symptomatic disease conditional on being infected, adjusted for age and exposure) were classified as “low immunity” and those with the lowest random effects classified as having relatively higher immunity, termed “moderate immunity” for this analysis to distinguish them from more immune adults.

*P. falciparum*-exposed individuals 1, 2, 141, 175, and 436 were anonymous blood donors at Mbale regional blood bank in Eastern Uganda. Plasma from anonymous *P. falciparum*-naïve U.S. donors was purchased from BioIVT. Samples from convalescent COVID-19 patients used in this study were received de-identified from the University of Texas Health San Antonio COVID-19 Repository. Systemic lupus erythematosus and rheumatoid arthritis patients were recruited at the University of Texas Health Science Center at San Antonio Rheumatology clinic.

### Generation of expression plasmids for PfARMA truncations

The plasmid encoding full-length PfARMA (PF3D7_1136200-bio, Addgene #47730) was modified to introduce a 6× His tag at the C-terminus. PfARMA fragments encoding IDR1, the folded domain, IDR2, IDR1 + the folded domain, and the folded domain + IDR2 were PCR amplified (**Table S16**) using site specific primers from this parent plasmid with overhangs encoding a NotI and AscI cut site added on the 5’ end and 3’ end, respectively. PCR amplified fragments and the parent plasmid were digested using NotI-HIFI (NEB #R3189S) and AscI (NEB #R0558S) according to manufacturer specifications. The digested plasmid was run on a 1% agarose gel with gel green for visualization. The plasmid backbone was cut from the 1% agarose gel and placed into a 1.5 mL Eppendorf tube. Agarose-dissolving buffer (Zymo #D4001-1-100) was added according to manufacturer specifications, and the gel was placed at 55°C until completely dissolved. Digested PCR fragments and plasmid backbone were purified using a Zymo DNA Clean & Concentrator kit (Zymo #D4004) according to manufacturer specifications and quantified using a NanoDrop One spectrophotometer (Thermo). Purified fragments and plasmid backbone were ligated using T4 DNA ligase (NEB #M0202S) according to the manufacturer specifications at a 3:1 ratio of insert to vector. After ligation, plasmids were transformed into MIX & GO! Competent DH5α cells (Zymo #T3007) according to the manufacturer specifications, plated onto agar plates containing ampicillin, and allowed to grow overnight. The following day, colonies were screened for inserts of the correct size using colony PCR and subsequently sequence-verified using whole plasmid sequencing (Plasmidsaurus).

### Recombinant protein expression and purification

Expi293F cells (Thermo #A14527) were maintained in a 37°C incubator with ≥80% relative humidity and 8% CO2 on an orbital shaker platform (125 rpm with a 19 mm shaking diameter). Cells were cultured in non-baffled polycarbonate flasks with a vented cap (Fisher #PBV12-5) and passaged every 3 to 4 days when the cell density was between 3 × 10^6^ and 5 × 10^6^ viable cells/mL. Absence of mycoplasma contamination was confirmed using the MycoAlerta Plus mycoplasma detection kit (Lonza #LT07705).

To produce C-terminally biotinylated proteins for B cell isolations, Expi293F cell were transfected with the relevant expression plasmid (strain 3D7 PfMSP1-bio, PfMSP3-bio, PfAMA1-bio, Pf41-bio, Pf113-bio, PfVFT-bio, and PfARMA-bio; Addgene #47709, #47731, #47741, #47739, #47729, #47790, and #47730, respectively) and a plasmid expressing secretedBirA-8his (Addgene #32408) at a 4:1 (w/w) ratio according to Thermo’s protocol. Biotin (Thermo #PI21336) was added to a final concentration of 100 μM immediately after adding the transfection mix. Culture supernatants were collected 5 – 7 days post-transfection by centrifuging the culture at 4,000 × g for 25 min. at RT. A 10 kDa cutoff Protein Concentrator PES (Thermo #88527) was used (5,000 × g at 4°C) to exchange culture medium containing free biotin for PBS (pH 7.2) (> 100,000 dilution) and to concentrate the protein to a final volume of 0.5 – 1 mL. The proteins were mixed with 6 – 12 volumes of PBS (pH 5.5) in a final volume of 6 mL and was subsequently loaded onto gravity flow columns (Thermo #29924) containing CaptAvidin agarose (Thermo #C21386) for purification. After three washes with PBS (pH 5.5) and five 6 mL elutions with PBS (pH 10.5), the elutions were pooled (30 mL) and the pH was immediately neutralized by adding 12 mL PBS (pH 5.5).

Plasmids encoding his-tagged full-length PfARMA and PfARMA truncations were transfected according to the manufacturer specifications and collected 6 – 7 days post-transfection. Cell suspensions were centrifuged at 4000 × g for 25 min at 4°C and supernatant was collected for protein purification. Protein purifications were preformed using a ÄKTA automated liquid chromatography system and HiTrap HP columns (Cytiva #29051021). Columns were washed with 5 column volumes (CV) of milliQ water and subsequently charged using 5 CV of 0.1 M NiSO_4_ solution. Column was then equilibrated with 5 CV of PBS. Subsequently, Expi293F supernatant containing recombinant protein was allowed to flow through the column using PBS. Columns were washed using 10 CV of PBS followed by 5 CV of 60 mM imidazole in PBS. After washing, bound protein was eluted in 10mL fractions using 300 mM imidazole in PBS. Following the elution step, fractions containing protein were buffer exchanged into PBS using a HiPrep 26/10 Desalt column (Cytiva #17508701).

After concentrating, the proteins were quantified using the Coomassie Plus (Bradford) Assay Kit (Thermo #23236) on a NanoDrop One spectrophotometer, according to the manufacturer’s instructions, diluted to 1 mg/mL, aliquoted, and stored at −70°C.

### Antigen-specific B cells isolations

On the day prior to the isolation of antigen-specific B cells, antigen tetramers were made by incubating biotinylated protein with streptavidin-PE (Cytek Biosciences #50-4317) or streptavidin-APC (Cytek Biosciences #20-4317) at a 6:1 molar ratio overnight at 4°C. Cryopreserved PBMCs from *P. falciparum*-exposed individuals were thawed at 37°C in a water bath and immediately mixed with pre-warmed thawing medium [IMDM Glutamax (Thermo #31980030) supplemented with 10% heat-inactivated fetal bovine serum (FBS) of US origin (Sigma #TMS-013-B) and 33 U/mL universal nuclease (Thermo #88700)] and then centrifuged for 5 min. at 250 × g at RT. Cells were resuspended in thawing medium and assessed for cell viability by adding 10 µL filtered 0.1% Erythrosin B in PBS to 10 µL of the cell suspension using a hemocytometer. Next, cells were centrifuged for 5 min. at 250 × g and RT and subsequently resuspended in isolation buffer (PBS supplemented with 2% heat-inactivated FBS and 1 mM EDTA) at 50 million live cells/mL and filtered through a 35 μm sterile filter cap (Corning #352235) to break apart any aggregated cells. B cells were then isolated using a MojoSort Human Pan B Cell Isolation Kit (BioLegend #480082) according to manufacturer’s instruction. The isolated B cells were washed with PBS and subsequently incubated with 1 µL LIVE/DEAD Fixable Aqua Dead Cell Stain Kit (Thermo #L34965) as indicated by the manufacturer. Next, B cells were washed with 2 mL cold PBS with 1% bovine serum albumin (BSA) (Sigma #A7979) and resuspended in 50 µL cold PBS with 1% BSA. B cells were then stained with a cocktail of *P. falciparum* antigen tetramers coupled to PE and APC (PfMSP1, PfAMA1, PfVFT, Pf113, PfMSP3, Pf41, and PfARMA, 25 nM each) for 30 min. in the dark on ice. The cells were then washed with 1 mL cold PBS with 1% BSA (5 min. at 250 × g at 4°C) and subsequently stained on ice for 30 min. with an antibody panel against B cell surface markers, consisting of Super Bright 645 anti-human CD19 (clone HIB19) 1:40 (Thermo #64-0199-42), Brilliant Violet 785 anti-human CD20 (clone 2H7) 1:50 (BioLegend #302355), PerCP-eFluor 710 anti-human CD21 (clone HB5) 1:40 (Thermo #46-0219-41), PE/cyanine7 anti-human CD27 (clone O323) 1:33 (BioLegend #302837), FITC anti-human IgG (clone G18-145) 1:40 (BD Biosciences #560952), PE/dazzle 594 anti-human IgD (clone IA6-2) 1:40 (BioLegend #348239), and eFluor 450 anti-human IgM (clone SA-DA4) 1:40 (Thermo #48-9998-41). UltraComp eBeads (Thermo #01222242) were used to prepare compensation controls for each fluorophore per manufacturer’s instructions. Prior to sorting, stained cells were washed with 3 mL cold PBS with 1% BSA (5 min. at 250 × g and 4°C) and subsequently filtered into a FACS tube with filter cap. Sorting was performed on a FACSAria II cell sorter. Lymphocytes were gated using forward and sideward scatter, followed by doublet exclusion and gating on live cells. *P. falciparum*-specific mature B cells (CD19^+^CD20^+^) were gated on tetramer binding (PE^+^APC^+^), and subsequently on IgG and IgM expression. IgM^+^ and IgG^+^ cells were sorted separately into 0.5 mL of IMDM/Glutamax/10% FBS in a 1.5 mL tube, diluted to a concentration of 1 cell per 100 µL, and plated into a 96-well plate to obtain approximately 1 cell per well (Corning #353072). One day prior to the sort, each well was seeded with 30,000 adherent, CD40L-expressing 3T3 cells in 100 µl IMDM/Glutamax/10% FBS containing 2× MycoZap Plus-PR (Lonza #VZA-2021), 100 ng/mL human IL-2 (GoldBio #1110-02-50), 100 ng/mL human IL-21 (GoldBio #1110-21-10), to promote expansion and differentiation of B cells into antibody-secreting cells [61,62]. After incubation at 37°C and 8% CO_2_ for two weeks, the wells were screened for the production of IgM or IgG by enzyme-linked immunosorbent assay (ELISA) and antigen-specificity was determined using Luminex assay to the select panel of *P. falciparum* antigens.

### Enzyme-linked immunosorbent assays

To detect IgG and IgM in B cell supernatant or plasma, 96-well ELISA plates (Corning #3361) were coated with either goat anti-human IgG (Sigma #I2136) or IgM (Sigma #I1636) antibody at a concentration of 4 and 8 μg/mL, respectively, diluted in PBS, at a total volume of 100 μL per well. After a one-hour incubation at 37°C or O/N at 4°C, each well was washed once using slowly running (approximately 900 mL/min.) deionized water. All subsequent washes were performed this way. Next, 150 μL blocking buffer (one-third Non-Animal Protein (NAP)-Blocker (G-Biosciences #786-190P) and two-thirds PBS) was added to each well to prevent non-specific binding. After one hour of incubation at 37°C, the wells were washed three times and 50 μL B cell culture supernatant diluted 1:1 in dilution buffer (1% NAP Blocker in PBS; total volume 100 μL) or plasma diluted 1:1,000,000 in dilution buffer was added per well. Plates were incubated for two hours at 37°C and washed five times. Then, either 100 μL 1:2500 diluted (1% NAP Blocker in PBS) HRP-conjugated anti-human IgG antibody (BioLegend #410902) or 1:5000 HRP-conjugated anti-human IgM antibody (Sigma #AP114P) was added to each well. After incubation for one hour at 37°C and three washes, HRP activity was detected using 50 μL TMB (Thermo #PI34024). Plates were incubated in the dark at RT and the oxidation reaction was stopped by adding 50 μL 0.18 M H2SO4 (Fisher #FLA300-212) per well when the negative controls (wells that received buffer when test wells received culture supernatant) started to color. Absorbance was measured at 450 nm using a BioTek Synergy H4 microplate reader. A human IgG (Sigma #I2511) or IgM (Sigma #I8260-1MG) standard curve (ten three-fold serial dilutions starting at 20 μg/mL) was used to quantify samples. For B cell supernatants, wells with values >27 ng/mL were considered positive. This cutoff was determined based on our observation that the amplification of heavy and light chain variable regions always failed from cultures with a lower concentration. For plasma IgG samples, the IgG concentration was interpolated from the standard curve using sigmoidal four-parameter logistic fitting in Graphpad Prism 10.

ELISAs to measure IgG reactivity to PfMSP1, PfARMA, or PfARMA fragments were performed as described above with the following modifications. Plates were coated with 50 μL in-house produced protein per well at a concentration of 2 μg/mL (0.1 μg/well). Coated plates were incubated for 1 hour at 37°C or overnight at 4°C and all subsequent incubations were done at RT instead of 37°C. To prevent non-specific binding, the wells were blocked with 200 μL PBS containing 0.1% tween-20 and 3% non-fat milk powder (SACO), which significantly increased specificity of the assay (compared to NAP blocker). After discarding the blocking buffer from the wells, the plates were not washed. Purified antibodies were tested at a final concentration of 2.5 μg/mL in 100 – 200 μL in PBS containing 0.1% tween-20 and 1% non-fat milk powder. The plates were washed six times prior to adding the detection antibody, and four times prior to adding TMB substrate.

### Luminex assays

Custom Luminex beads were generated by coupling 10 pmol of each protein (PfMSP1, PfAMA1, PfMSP3, Pf41, Pf113, PfVFT, PfARMA, PfARMA IDR1, PfARMA folded domain, and PfARMA IDR2) per 1 × 10^6^ MagPlex microspheres (Luminex #MC10025-ID) using the Luminex protein coupling kit (#40-50016) per manufacturer’s instructions. The coupled beads were stored at 4°C in the dark. Buffer A (PBS with 0.05% Tween 20 (Fisher #BP337), 0.5% BSA (Sigma #A7979), 0.02% sodium azide) and Buffer B (0.05% Tween-20, 0.5% BSA, 0.02% sodium azide, 0.1% casein (Sigma #C7078), 0.5% PVA (Sigma #P8136) and 0.5% PVP (Sigma #PVP360), 15 µg/mL *E. coli* lysate) were prepared ahead of time and also stored at 4°C. For buffer B, the chemicals were allowed to dissolve O/N, and *E. coli* lysate (MCLAB #ECCL-100) was added the next day, followed by centrifugation at 10,000 × g for 10 min. to clear the buffer. All assay steps were done at RT and the beads were protected from light using aluminum foil. Coupled beads were pooled, resuspended in buffer A and plated at 1000 beads per well for each protein in a black, flat-bottom 96 well plate (Bio-Rad #171025001). The beads were washed once. All washes were done with 100 µL PBST (PBS with 0.05% Tween 20) using a handheld magnetic washer (Bio-Rad #171020100). The incubation time on the magnet was always 2 min. Next, the beads were incubated with 50 µL of human serum (diluted 1:200 using buffer B) or B cell culture supernatant (diluted 1:1 with buffer B) for 30 min. with constant agitation (500 rpm, 2.5 mm orbital diameter). After three washes, 50 µL secondary antibody diluted in buffer A (PE anti-human IgG (1:200 dilution; Jackson ImmunoResearch #109-116-098) or PE anti-human IgM (1:200 dilution; Jackson ImmunoResearch #109-116-129)) was added per well. After 30 min. incubation with constant agitation, the beads were washed three times and subsequently incubated in 50 µL buffer A for 30 min. with constant agitation. After one final wash, the beads were resuspended in 100 µL PBS and fluorescence intensity was measured using a calibrated and validated Bio-Rad Bio-Plex 200 machine. Samples were analyzed in duplicate and were excluded from the analysis when the two measurements differed by more than two-fold. To determine the cutoff for IgG seropositivity to full-length PfARMA and IDR1 in samples from *P. falciparum*-naïve individuals, a Gaussian mixed model was used to detect a negative and a positive subset among the samples. The model was created in R (v4.4.1) using the package mclust (v6.1.2) [63].

### Amplification of antibody heavy and light chain variable regions

PfARMA-specific, monoclonal B cell cultures were collected by centrifugation (5 min. at 250 × g and RT), resuspended in 50 μL Tri-Reagent (Zymo #R2050-1-200) and stored at −70°C. Heavy and light chain variable regions were amplified from CIDRα1-specific B cells after cDNA synthesis and a series of PCR reactions as described [45]. All primer sequences can be found in **Table S17**. RNA was isolated using Zymo’s Direct-zol RNA Microprep kit (#R2060), eluted in 15 μL nuclease-free water and then mixed with 0.7 μL reverse primer (10 μM, 200 mM final concentration (f/c) in 35 μL PCR reaction volume) specific for the IgG or IgM heavy chain (primer #7 and #297) plus 0.7 μL light chain specific reverse primers (10 μM): #108 and #109), and incubated for two minutes at 65°C. Single stranded cDNA was synthesized immediately by adding 7 μL First-Strand buffer (f/c 1×), 7 μL DTT (20 mM, f/c 4 mM), 0.7 μL dNTPs (10 mM each, f/c 200 μM each, Sigma #DNTP-10), 1.75 μL RNase OUT (40 U/μL, f/c 2 U/μL, Thermo #10777019), 0.7 μL template switch oligo (TSO) (10 μM, f/c 200 nM, IDT DNA; #110, **Table S17**), 0.7 μL SMARTScribe reverse transcriptase (100 U/μL, f/c 2 U/μL, Takara Bio #639537), nuclease-free water till 35 μL, and subsequent incubation at 42°C for 2 hours. The TSO was designed with two modified bases, iso-dC and iso-dG, at the 5’ end to prevent TSO concatemerization, and three riboguanosines at the 3’ end for increased binding affinity to the appended deoxycytidines (property of the Takara reverse transcriptase) [64,65]. The single-stranded cDNA was immediately purified using Zymo’s RNA Clean & Concentrator kit (#R1016) using Zymo’s appended protocol to purify fragments >200 nucleotides and was eluted in 10 μL elution buffer. This critical clean-up step ensured that any unused TSO was removed, preventing it from inhibiting the subsequent PCR reactions by serving as template for the forward primer. Immediately after, heavy and light chain variable regions were amplified by PCR in one reaction mix using 8.5 μL purified cDNA, 10 μL 2× AccuStart II PCR SuperMix (QuantaBio #95137), 0.9 μL 10 μM forward primer #106 (f/c 0.45 μM, **Table S17**), and 0.2 μL of the reverse primers (10 μM) used to synthesize the cDNA (#7, #297, #108, and #109, each at f/c 0.1 μM). Cycling conditions were 94°C for 3 min., 35 cycles of 30 sec. at 94°C, 30 sec. at 55°C and 35 sec. at 72°C, followed by 5 min. at 72°C. A second, nested amplification was required to obtain enough amplicon DNA, and was done separately for heavy chain, kappa light chain, and lambda light chain variable regions, using AccuStart II PCR SuperMix, and 2 μL of the first, unpurified PCR as template in a total reaction volume of 20 μL. Mixes of primers as described by Hua-Xin Liao *et al.* [66] were used for this second PCR, with a final concentration of 0.1 μM for each individual primer. Reverse primer #67 was added for the heavy chain variable region PCR to allow for amplification of variable regions originating from IgG_2_, IgG_3_ and IgG_4_ mRNA, in addition to #30 which is specific for IgG_1_. Cycling conditions were as described above, except for the extension step (shortened to 30 sec.) and the annealing step, which was 30 sec. at 60°C for the IgG_1_ heavy chain variable region, 30 sec. at 63°C for the IgM heavy chain variable region, and 30 sec. at 50°C for the light chain variable regions. For sequencing of the variable regions and antibody expression, linear IgG expression cassettes were generated as described [45,66]. Variable region sequences were analyzed using the International Immunological Information System (IMGT) gene database and the V-QUEST sequence alignment tool [67] using default settings to identify V(D)J gene usage and amino acid substitutions.

### Generation of antibody expression plasmids

Antibody variable regions were cloned into expression plasmids from Invivogen (#pfusess-hchg1, #pfuse2ss-hclk, #pfuse2ss-hcll2). The variable heavy and light chain regions were amplified from the linear expression cassettes (2 µL at 1 ng/µL) using 10 µl NEB Q5 Hot Start HiFi PCR master mix (#M0494S), 6 µL nuclease-free water and 1 µL sequence-specific F and R primer (10 µM, f/c 500 nM, **Table S16**) that were based on the results of analysis using IMGT/VQUEST [67]. These primers introduced restriction sites (EcoRI & NheI for hchg1, EcoRI & BsiWII for hclk, and EcoRI & AvrII for hcll2). Every plasmid was Sanger sequence-verified prior to using it as expression vector.

### Antibody expression and purification

Heavy and light chain antibody expression plasmids were used at a molar ratio of 1:2 to transfect 5 mL cultures. The antibodies were purified from the culture supernatant 4 – 6 days later using protein G magnetic beads (Promega #G7472). Purified antibodies and antibody elution buffer [5 parts elution buffer (100 µM glycine-HCl, pH 2.7) and 1 part neutralization buffer (2 M Tris buffer, pH 7.5)] were buffer exchanged to PBS using 100 kDa cutoff Protein Concentrators (Thermo #88523). The samples were diluted > 50,000 × in PBS by repeated centrifugation at 4,000 × g and 4°C. Purified antibodies were quantified using the Coomassie Plus (Bradford) Assay Kit (Thermo #23236) on a NanoDrop One spectrophotometer, according to the manufacturer’s instructions, and visualized on SDS-PAGE gel with a standard amount of BSA to evaluate protein size and purity.

### Dot-blot assay

Two µL of purified protein (100 ng/µL) or 2 µL of unpurified protein was spotted on a 0.45 µm nitrocellulose membrane (Thermo # PI88014). Membranes were allowed to dry at RT for 45 min. All subsequent incubation steps were performed at RT on a platform rocker. After drying, membranes were blocked for non-specific interactions with 5% non-animal protein (NAP, G-Biosciences #786-190P) in 0.05% Tween-20 in 1× PBS (0.05% PBST) for 1 hour. After the blocking step, membranes were incubated with 1 µg/mL of primary antibody in 5% NAP in 0.05% PBST for 30 min. After primary antibody incubation, membranes were washed three times with 0.05% PBST for 5 min. Subsequently, membranes were incubated with anti-IgG HRP-conjugated secondary antibody (1:1000 dilution, Biolegend #490902) in 0.05% PBST for 30 min. Membranes were then washed three times with 0.05% PBST for 5 min. Afterwards, membranes were rinsed twice with deionized H_2_O and incubated with 1-Step Ultra TMB Blotting Solution (ThermoFisher #37574) for 5 min. Membranes were washed with deionized H_2_O and imaged on a BioRad ChemiDoc MP Imaging System (BioRad #12003154).

### Biolayer interferometry

hmAb binding was assessed using biolayer interferometry on an Octet 96 Red (ForteBio) or RH16 (Sartorius) using AHC biosensors (Sartorius). Purified IgGs were diluted to 20 μg/mL in kinetics buffer (1× PBS, 0.01% Tween 20, 0.01% BSA and 0.005% NaN_3_ (pH 7.4)). IgGs were loaded onto the biosensor for 120 s. After loading, biosensors were placed in kinetics buffer to 60 s for a baseline reading. Biosensors were then immersed in analyte at a concentration of 1 μM in kinetics buffer for 300 s in the association phase, followed by 300 s in the dissociation phase in kinetics buffer. The background signal from a biosensor loaded with IgG but with no analyte was subtracted from each loaded biosensor.

To facilitate expression of the IDR1 peptide 220-269, a fusion protein using maltose binding protein (MBP) as a carrier was made. A plasmid containing an N-terminal his-tag, mammalianized MBP [68], HRV3C cleavage site and C-terminal ARMA 220-269 was synthesized by Twist Bioscience in the pTwist-CMV plasmid. The fusion protein was expressed in HEK293E cells (National Research Council, Canada) using PEI as a transfection reagent. Cultures were harvested after 7 days, and protein were purified from culture supernatant using His60 Ni Superflow resin (Takara Bio #635660) and eluted from the column using a buffer of 50 mM Tris, 300 mM NaCl (pH 7.5) with 150 mM imidazole. Fusion protein was further purified using a Superdex 200 16/600 size-exclusion column (Cytiva #28-9893-35) using an AKTApure system (GE Biosciences).

### Parasite culture

*P. falciparum* strain 3D7 parasites were cultured [69] in human O^+^ erythrocytes from local blood donors at 3 – 10% parasitemia in complete culture medium (5% hematocrit). Complete culture medium consisted of RPMI 1640 medium (Gibco #32404014) supplemented with gentamicin (45 µg/mL final concentration; Gibco #15710064), HEPES (40 mM; Fisher #BP3101), NaHCO_3_ (1.9 mg/mL; Sigma #SX03201), NaOH (2.7 mM; Fisher #SS266-1), hypoxanthine (17 µg/mL; Alfa Aesar #A11481-06), L-glutamine (2.1 mM; Corning #25005Cl), D-glucose (2.1 mg/mL; Fisher #D16-1), and 10% heat-inactivated human AB^+^ serum (Valley Biomedical #HP1022). Parasites were cultured at 37°C in an atmosphere of 5% O_2_, 5% CO_2_, and 90% N_2_. Before use in cultures, 12.5 mL packed erythrocytes were washed twice with 10 mL cold incomplete medium (complete culture medium without human serum) and pelleted between each wash by centrifugation at 500 × g for 8 min. at 4°C (max. acceleration and slow break). Washed erythrocytes were resuspended in 1 volume of complete medium to 50% hematocrit and stored at 4°C.

Parasites were synchronized to the ring stage by treatment with 5% D-sorbitol [70] (Fisher #S459-500). Cultures containing high percentages of ring-stage parasites were centrifuged at 250 × g for 5 min. at RT. Pelleted erythrocytes were resuspended in 10 volumes of 5% D-sorbitol in MQ water, vortexed for 30 sec. and incubated for 8 min. at 37°C. The cells were vortexed another 15 sec., washed with 5 volumes of complete culture medium (250 × *g* for 5 min. at RT), and resuspended in complete culture medium at 5% hematocrit and cultured as described above.

### Growth inhibition assay

*P. falciparum* isolate 3D7 parasites were pre-synchronized at the ring stage with a 5% D-sorbitol (Fisher #S459-500) treatment as described above, followed four days later by two additional 5% D-sorbitol treatments 14 hours apart [70]. At the late trophozoite/early schizont stage (24 hours after the third D-sorbitol treatment), parasitemia was determined by inspection of a Giemsa-stained blood smear. The smear was also used to confirm correct parasite staging. Immediately after, 40 µL of each antibody (2 mg/mL in PBS) was added per well of a 96-well half-area microplate (Corning #3696). Two-fold serial dilutions were made by pipetting 20 µL of the antibody in the first row and adding this to the second row containing 20 µL PBS. After mixing by pipetting, 20 µL was removed from this row and pipetted into the next row containing 20 µL PBS, for a total of 4 concentrations (2 mg/mL – 0.25 mg/mL). A monoclonal antibody specific for PfAMA1 was used as a positive control (BEI #MRA-481A). An PfAMA1-specific monoclonal antibody with no reported growth inhibitory activity was used as a negative control (BEI #MRA-480A). Wells with PBS were included as a reference for parasite growth in the absence of antibody. An equal volume of parasite culture (1% parasitemia and 2% hematocrit) was then added to all wells containing antibody solutions or PBS. Uninfected erythrocytes (2% hematocrit) were used to determine the background signal. All conditions were performed in triplicate. Empty wells and inter-well spaces were filled with PBS to minimize evaporation. The plate was then incubated at standard parasite culture conditions (described above) for 48 hours. Next, erythrocytes were washed by adding 120 µL cold PBS to each well, and centrifuging the plate for 5 min. at 1,400 × g at 4°C. After removing 120 µL from each well, this wash step was repeated once. The plate was then placed on a shaker at 1000 rpm (2.5 mm orbital diameter) for 30 sec. or until the pellets had been completely resuspended. To perform readout of the assay, 120 µL of LDH substrate (0.1 M Tris-HCl, pH 8.0, 50 mM sodium L-lactate, 0.25% (v/v) Triton X-100, 0.5 mg/mL nitro blue tetrazoline (Sigma #N5514), 50 µg/mL 3-Acetylpyridine Adenine Dinucleotide (Sigma #A5251), 1 U/mL Diaphorase (Sigma #D5540)) was added per well, and the plate was spun for 1 min. at 1,800 × g at RT. After 10 min., the plate was placed in a BioTek Synergy H4 plate reader, shaken for 15 sec. and absorbance at 650 nm was read. If the absorbance in the wells with infected erythrocytes incubated with PBS only was below 0.4, the plate was measured again at 5-min. intervals until the absorbance in these wells was between 0.4 and 0.6. The average background value was subtracted from the signal of the wells with infected cells. Percent growth inhibition was expressed as the reduction in signal in wells incubated with antibody as compared to the negative control.

### PhIP-seq assay

The *P. falciparum* T7 phage has been described previously [27], and samples were processed with some modifications to the prior procedure. Plasma was diluted to 0.1× in CSF buffer (20 mM HEPES, pH7.3, 0.02% NaN_3_, 20% glycerol in PBS, pH7.4), and monoclonal antibodies were diluted to 50 ng/μL in CSF buffer. Five microliters of each sample were then added to wells containing 500 µL of phage suspension (∼2.5 × 10^10^ PFU per well), with beads only control wells receiving CSF only. Plates were sealed with a silicone mat and rotated end-over-end overnight at 4°C. For immunoprecipitation, 15 μL of suspension from Sera-Mag SpeedBeads Protein A/G Magnetic Particles (Cytiva #17152104010350) was washed three times with a 30 μL volume of Tris-Buffered Saline (TBS) + 0.5% Triton X-100 before dispensing to each sample well. Plates were sealed and rotated for an additional hour at 4°C. Protein A/G beads with bound immunoglobulin plus phage were washed six times in a combined wash volume of 2.5 mL of TBS + 0.5% Triton using a plate magnet (Alpaqua) and 96-channel pipettor (Rainin). Beads were then washed in 200 μL PBS without Ca^2+^ or Mg^2+^ (Corning #21-040-CM), resuspended in PBS, transferred to a PCR plate, followed by removal of all remaining PBS. The PCR plate containing the bead pellets was sealed with a foil seal and stored at −80°C.

The sequencing libraries were prepared by direct PCR of phage bound to the washed beads. Sample plates were thawed on ice. For PCR1, each well containing approximately 2 μL of washed, pelleted beads received 23 µL of a mixture of 12.5 μL of NEBNext Ultra II Q5 Master Mix (NEB #M0544X), 0.5 μL each of the 10 μM forward and reverse primer pools [27], and nuclease-free water for a final 25 μL reaction volume. Initial lysis and denaturing was at 98°C for 2 minutes, followed by 28 cycles of 98°C for 5 seconds, 70°C for 20 seconds, and 72°C for 15 seconds with a final 2-minute extension at 72°C. For PCR2 (dual-indexing), 0.5 μL of the PCR1 product was mixed into a final 12.5 μL reaction volume with the same 2× Q5 Master Mix along with 1 µL of 5 μM each of custom 12-bp TruSeq-compatible barcodes [27]. The PCR2 reaction was amplified for 5 cycles as above for PCR but with 30 seconds of initial denaturation. Each indexed sample (5 μL) was pooled and bead cleaned using in-house SPRI beads. PhiX (10%) was added to the pooled sequencing library before sequencing on a single lane of a NovaSeqX 10B PE150 flow cell (MedGenome, Foster City CA).

Reads were demultiplexed and assigned to peptide sequences using kallisto [71] to obtain peptide counts. The function runEdgeR() with de.method = “glmQLF” from the R package BEER [72] was used to estimate each peptide’s log2 fold change declaring beads-only sample wells (n = 14) as the ‘beads’ background binding population. P values were adjusted for multiple hypothesis testing using the Benjamini-Hochberg procedure.

### Immunofluorescence assay

For immunofluorescence staining, a thin blood smear was made on a microscopy slide using 1 µL of synchronized, late-stage *P. falciparum* parasite cultures. The blood smears were allowed to dry for 30 seconds at RT and subsequently fixed using 1 mL of 4% paraformaldehyde (Electron Microscopy #15710). Slides were incubated with 4% paraformaldehyde for 30 min. at RT and were then washed three times with 1 mL 1× PBS and permeabilized using 0.1% Triton-X (Fisher # BP151) in 1× PBS. Slides were incubated with 0.1% Triton-X for 30 min. at RT and subsequently washed three times using PBS. Next, blocking buffer (2% BSA, 0.05% Tween-20, 100 mM Glycine, 3 mM EDTA and 150 mM NaCl in 1× PBS) was added and incubated for one hour at RT. Slides were washed three times and primary antibody was added at 5 µg/mL in 500 µL blocking buffer. Slides were incubated with 5 µg/mL primary antibody (human anti-PfARMA 5.314; mouse anti-PfRON4, WEHI Antibody Facility #6A10; mouse anti-PfAMA1, BEI #MRA-481A; mouse anti-PfMSP1, BEI #MRA-880A; mouse anti-PfRAP1, BEI #MRA-833; mouse anti-EBA-175, BEI #MRA-711A; mouse anti-RESA, WEHI Antibody Facility #28/2) for 1 hour at RT and subsequently washed three times using 1× PBS. Secondary antibodies (goat anti-human IgG – Alexa Fluor 647, Thermo #21445 and goat anti-mouse IgG – Alexa Fluor 555, Thermo #A32727) were diluted 1:1000 (1.5 µg/mL) in blocking buffer and then added to the smear to incubate for one hour in the dark. For the positive control, samples were stained with primary anti-RAP1 antibody and secondary antibodies goat anti-mouse IgG – Alexa Fluor 555 and goat anti-mouse IgG – Alexa Fluor 647 (Thermo # A32728). Samples were again washed three times using PBS in the dark and then allowed to air-dry for one hour in the dark. Slides were mounted using 10 µL ProLong Glass mounting medium containing NucBlue Stain (Thermo #P36985) and sealed with a cover slip. Samples were imaged using a Zeiss Axio Imager Z1 with Apotome using Zen Blue software.

### Colocalization analysis

Pearson’s correlations and Mander’s coefficients were calculated using the BIOP-JACoP plugin on the Fiji image analysis software (v2.9.0). Five images stained with anti-PfARMA antibody and antibody targeting the previously localized antigen (PfMSP1, PfAMA1, PfEBA-175, PfRON4, PfRAP1, and PfRESA) were selected for analysis. Five images stained with anti-PfRAP1 and two secondary antibodies targeting the anti-PfRAP1 antibody were used as positive controls. To minimize the non-parasite space analyzed, images were cropped to only include the parasite of interest. The fluorescent signal associated with PfARMA staining was assigned to one channel and the signal associated with the previously localized antigen (PfMSP1, PfAMA1, PfEBA-175, PfRON4, PfRAP1, and PfRESA) was assigned to the other channel. For the positive control, one fluorescent signal was assigned to channel A and the second was assigned to channel B.

### Plasma IgG purification

One mL of plasma was thawed at RT and diluted 1:1 with 1 mL of sterile 1× PBS (Gibco #20017-027). Diluted plasma was subsequently centrifuged at 16,000 × g for 3 min. at RT to pellet aggregates and the supernatant was transferred to a 3 mL syringe (BD #309657). A 5 mL polypropylene column (Thermo #29922) was placed on a ring stand and a frit was inserted into the bottom of the column. A total of 1.5 mL 50% Protein G Plus slurry (Thermo #22852) was transferred to the column and 8 mL PBS was added to the column to neutralize the resin. A second frit was added to the column and the PBS was allowed to flow out of the column. A 45 µm syringe filter (Fisher #09-720-4) was placed on the syringe containing the diluted plasma and contents were transferred to the column. The syringe was rinsed with 1 mL PBS and this volume was also added to the column. The flowthrough (2 mL diluted plasma and 1 mL PBS) was collected in a 15 mL conical tube and ran over the column an additional 2 times (3 times total, flowthrough collected each time). To push out the remaining plasma, 2 mL PBS was added to the column and collected in the 15 mL conical tube containing the diluted plasma flowthrough. The 15 mL conical tube containing the plasma flowthrough was stored at −80°C. Next, the column was washed with 30 mL of PBS. During this time, six 2 mL Eppendorf tubes were prepared for collecting elution fractions by adding 200 µL 1 M Tris-HCl (pH 8.0). After washing, 1 column volume (0.75 mL) of elution buffer (0.1 M glycine, pH 2.7) was added to the column to push out the remaining PBS and collected in the first elution tube. Next, 9 mL of elution buffer was added to the column and 1.8 mL fractions were collected in the 5 remaining elution tubes. Tubes were mixed to neutralize the elution buffer. To ensure complete neutralization, the pH was assessed by adding 0.5 µL elution to a pH indicator strip (Fisher #13-640-517). Elution fractions were quantified for protein and fractions containing >0.15 mg/mL were pooled. Pooled fractions were concentrated and buffer exchanged into 1× PBS using a 5 to 20 mL 10k MWCO Pierce Protein Concentrator (Thermo #88527). First, pooled eluted IgG (9 mL) was added to the protein concentrator and topped off with PBS to a final volume of 20 mL. Samples were centrifuged at 4000 × g until the remaining volume was <500 µL. Then, 19.5 mL PBS was added to the concentrator and samples were centrifuged at 4000 × g until the remaining volume was <500 µL. This was repeated an additional 2 times. After concentrating and buffer exchanging, the purified plasma IgG was stored at 4°C for autoantibody or antigen-specific purification the following day.

### Autoantibody purification

To prepare for preparing human cell lysate, one tablet of cOmplete protease inhibitor (PI) cocktail (Roche #04693116001) was resuspended in 5 mL of cold PBS to generate a 10× PI cocktail. The 10× PI cocktail was aliquoted and stored at −80°C. This 10× PI cocktail was then diluted in cold PBS to generate a 2× PI cocktail working solution. A culture of Expi293F cells was centrifuged at 125 × g for 5 min. The cells were resuspended in 2× PI cocktail at a concentration of 6.25 × 10^4^ cells/µL and divided into 400 µL aliquots in 1.5 mL tubes (25 × 10^6^ cells total per tube). Tubes were snap-frozen in a dry ice/ ethanol slurry and stored at −80°C.

Expi293F cells were physically lysed through high frequency sonification and repeated freeze-thaw cycles to extract native human protein. Tubes containing 25 × 10^6^ Expi293F cells in 2× PI cocktail were thawed on ice and processed immediately once thawed. Sonication was done using a Branson Model SFX using an 11 mm-wide probe at 50% output power using a 10 second on, 10 second off cycle for a total of 2 minutes. The sonicator probe was placed 1 inch deep into a 1 L beaker containing ice and water and cells were sonicated by touching the bottom of the 1.5 mL tube to the sonicator probe in the ice and water mixture. Immediately following the completion of the sonication cycle, the 1.5 mL tubes containing the Expi293F cells were placed in a tube rack and frozen in a dry ice/ethanol slurry. Once frozen, cells were allowed to thaw at room temperature with intermittent agitation to ensure even thawing. After cells were complete thawed, freeze/ thaw was repeated for a total of 2 freeze/thaw cycles. Once cells were completely thawed from the second freeze/thaw cycle, the cell lysate was centrifuged at 264 × g for 5 min. After centrifugation, supernatants containing total human protein were pooled and quantified via Bradford assay. Expi293F cell protein was stored in aliquots of 12.5 mg total protein (concentration 10 mg/mL – 35 mg/mL) at −80°C.

Expi293F protein was coupled to pre-activated NHS-ester Sepharose for the isolation of human autoantibodies from plasma-isolated IgG. A total of 25 mg of Expi293F protein was thawed for each affinity column generated. Once thawed, protein aliquots were centrifuged at 125 × g for 5 min. and the supernatant was transferred to a new 1.5 mL tube. A 10 mL polypropylene column (Thermo #29924) was placed on a ring stand for each sample to be purified and a frit was placed into the bottom of the column. Next, 6 mL of 50% NHS-activated Sepharose slurry in 100% isopropanol (3 mL sepharose resin; Cytiva #17090601) was placed into each of the columns and the buffer was allowed to drain. Next, 15 mL of ice cold 1 mM HCl was added into each column and allowed to drain, followed by capping the bottom of the column. The NHS-activated Sepharose was resuspended in 4.5 mL of Expi293F protein diluted to a concentration of 5.5 mg/mL in coupling buffer (0.2 N NaHCO_3_, 0.5 M NaCl, pH 8.3). The columns were sealed and placed on a nutator mixer at 4°C overnight.

The following day, the column was placed back on a ring stand and the buffer was allowed to drain. Next, 5 m of blocking buffer (0.5 M ethanolamine, 0.5 M NaCl, pH 8.3) was added to the column and allowed to drain. The bottom of the column was capped and 10 mL of blocking buffer was added to the column. The column was sealed and placed on a nutator mixer at room temperature for 3 hours. Following the 3-hour incubation, the column was centrifuged inside a 50 mL conical tube at 200 × g for 10 min. After centrifugation, the top frit was placed into the column and the blocking buffer was subsequently drained. The column was then washed twice with 2 mL of blocking buffer, washed twice with 2 mL of wash buffer (0.1 M acetic acid, 0.5 M NaCl, pH 4), and washed twice with 2 mL of blocking buffer. The column was capped and 2 mL of blocking was added to the column. The column was left at RT in blocking buffer for 15 min. and then allowed to drain. Subsequently, the column was washed twice with 2 mL of wash buffer, twice with 2 mL of blocking buffer, and twice with 2 mL of wash buffer. Next, 5 mL of binding buffer (50 mM Na_2_HPO_4_, pH 7.0) was added to the column and allowed to drain. Finally, 5 mL of 20% ethanol was added to the column, 3 mL was allowed to drain before capping the column and storing at 4°C overnight.

On day 3, binding buffer (50 mM Na2HPO4, pH 7.0) and elution buffer (0.1 M glycine, pH 2.7) were brought to room temperature. Next, 3 mL plasma purified IgG was diluted 1:1 in 3 mL binding buffer. The column containing NHS-activated Sepharose coupled to Expi293F protein was placed on a ring stand and washed with 5 mL of binding buffer. Next, the column was washed with 5 mL of elution buffer. Finally, the column was equilibrated by adding 5 mL of binding buffer and allowing it to drain. Columns were capped and 6 mL of purified plasma IgG diluted in binding buffer was added and allowed to incubate on a nutator mixer for 1 hour at RT. Following the 1-hour incubation, the column was placed back on a ring stand and the unbound plasma IgG was allowed to drain. Unbound plasma IgG was collected and stored at 4°C for concentration and buffer exchange following the elution step. The column was washed twice with 10 mL of binding buffer and allowed to completely flow through. Next, 5 mL of elution buffer was added to the column and collected in a 15 mL conical tube containing 300 µL 1 M Tris-HCl pH 8.0. Elution sample was mixed to ensure complete neutralization. Autoantibody fraction and unbound plasma IgG were concentrated and buffer exchanged into 1× PBS using a 6 mL 10k MWCO Pierce Protein Concentrator (Thermo #88516). After samples were added, the Pierce Protein Concentrator was centrifuged at 4000 × g until the remaining volume was <200 µL. Then, 5.8 mL PBS was added to the column and samples were centrifuged at 4000 × g until the remaining volume was <200 µL. This was repeated an additional 3 times. After concentration and buffer exchanging, the purified autoantibody and unbound plasma IgG samples were quantified via protein Bradford assay and stored at −20°C.

### ARMA-specific antibody purification from plasma

The isolation of ARMA-specific antibodies from plasma IgG was performed as described above for the isolation of autoantibodies, with the following modifications. For each affinity column, 0.5 mg of recombinant PfARMA protein was thawed and 2 mL of 50% NHS-activated Sepharose slurry in 100% isopropanol (1 mL sepharose resin) was used. The NHS-activated Sepharose was resuspended in 2 mL of recombinant PfARMA protein diluted to a concentration of 0.25 mg/mL in coupling buffer.

### Polyreactivity ELISA

High-protein-binding plates (Costar #3361) were coated overnight at 4°C with the following antigens prepared in 1× PBS: 10 µg/mL calf thymus double-stranded DNA (Sigma #4522), 10 µg/mL calf thymus single-stranded DNA (Sigma #D8899), 5 µg/mL keyhole limpet hemocyanin (Thermo #77600), 5 µg/mL cyan fluorescent charged-variant proteins (produced in-house) [73], 10 µg/mL Hep-2 cell lysate (produced in-house), 5 µg/mL 2,4-dinitrophenol-bovine serum albumin (Thermo #A23018). Cardiolipin (Sigma-Aldrich #C0563) was dissolved at 50 µg/mL in 100% ethanol and coated under the same conditions. Plates were washed once with PBS containing 0.05% Tween-20 (PBST), or PBS for cardiolipin-coated wells, and blocked for 2 h at room temperature with 50 µL of blocking buffer (0.5% [w/v] BSA in PBST, or 10% [v/v] FBS in PBS for cardiolipin). After three additional washes, wells were incubated with primary antibody (100 nM in blocking buffer) in duplicate. Each antibody was also incubated at 100 nM in duplicate antigen-free control wells (BSA or FBS, as appropriate). As positive controls, we included anti-DNA mAb 3H9 [74] and mAb 4A5 that binds the S2 subunit of SARS-CoV-2 [75] and shows broad polyreactivity (Dr. Jennifer Maynard, personal communication). As a negative control, anti-CD20 mAb rituximab was used. Following three washes, plates were incubated for 1 h with goat anti-human IgG-HRP (1:5,000; Sigma-Aldrich #A0293). After three final washes, 50 µL of 1-Step Ultra TMB substrate (Thermo #34028) was added and the reaction was quenched after 15 min of shaking with 50 µL of 2 M H_2_SO_4_. Absorbance at 450 nm was measured using a Synergy H1 microplate reader.

### Statistics

Statistical analysis was performed using GraphPad PRISM 10 software. For the analysis of continuous values with non-linear differences (MFI and OD) between more than two groups, data were analyzed using the Kruskal-Wallis test, followed by comparisons between all pairs of groups using Dunn’s post-hoc test. For the comparison of continuous values with linear differences between two groups, data were analyzed using a student’s t-test. P values < 0.05 after correction for multiple testing were considered statistically significant.

## FUNDING

This work was funded by the National Institutes of Health/National Institute of Allergy and Infectious Diseases (R01 AI153425 to EMB). Collection of clinical samples was supported by the National Institutes of Health/National Institute of Allergy and Infectious Diseases (U19 AI150741 to BG and U19 AI089674 which funds the PRISM cohorts) and The Max and Minnie Tomerlin Voelcker Fund (Young Investigator Award to EMB). RG is enrolled in the South Texas Medical Scientist Training Program (STX-MSTP), supported by the NIH training grant T32GM145432 and fellowship award F30 AI176697. Data were generated in the Flow Cytometry Shared Resource at the University of Texas San Antonio Health Science Center, which is supported by a grant from the National Cancer Institute (P30 CA054174-20) to the Mays Cancer Center, a grant from the Cancer Prevention and Research Institute of Texas (CPRIT; RP210126), a grant from the National Institutes of Health (S10 OD030432), and the Office of the Vice President for Research at the University of Texas San Antonio Health Science Center.

## ACKNOWLEDGEMENTS

We thank Drs. Zhenming Xu and Maria Fernandez for sharing plasma samples of SLE patients and Dr. Augustin Escalante for the recruitment of RA patients. The 3T3-msCD40L cell line was a gift from M. Connors. Plasmids encoding *P. falciparum* 3D7 MSP1-bio, MSP3-bio, AMA1-bio, P41-bio, Pf113-bio, PF3D7_0606800-bio, PF3D7_1136200-bio, and birA were a kind gift from Dr. Gavin Wright (Wellcome Sanger Institute; Addgene plasmids #47709, 47731, 47741, 47739, 47729, 47790, 47730, and 32408). mAb 4A5 was a kind gift from Dr. Jennifer Maynard (University of Texas at Austin). The following reagents were obtained through BEI Resources, NIAID, NIH: *Plasmodium falciparum*, Strain 3D7, MRA-102, contributed by Dr. Daniel J. Carucci; anti-*Plasmodium falciparum* apical membrane antigen 1 (AMA1) monoclonal antibodies N3-1D7 (NRA-480A) and N4-1F6 (MRA-481A) (produced *in vitro*), contributed by Dr. Carole A. Long; anti-*Plasmodium falciparum* merozoite surface protein 1 (MSP1) monoclonal antibody, clone 5D5/47 (produced *in vitro*), MRA-880A, and hybridoma 7H8/50 anti-*Plasmodium falciparum* rhoptry-associated protein 1 (RAP1), MRA-833, contributed by Allan Saul; and anti-*Plasmodium falciparum* erythrocyte binding antigen-175 RII monoclonal antibody, clone R217 (produced *in vitro*), MRA-711A, contributed by B. Kim Lee Sim and NIAID/NIH.

## AUTHOR CONTRIBUTIONS

Conceptualization: Rolando Garza, Evelien M. Bunnik

Formal analysis: Rolando Garza, Sebastiaan Bol, Evelien M. Bunnik

Funding acquisition: Rolando Garza, Bryan Greenhouse, Evelien M. Bunnik

Investigation: Rolando Garza, Jeffrey Marchioni, Jared Honeycutt, Nicholas K. Hurlburt, Caroline Torres, Anakaren Garcia, Eva Loranc, Emily Yemington, Dalton Towers, Sebastiaan Bol, Evelien M. Bunnik

Project administration: Evelien M. Bunnik

Resources: Isaac Ssewanyana, Prasanna Jagannathan, Bryan Greenhouse

Supervision: Jason Lavinder, Marie Pancera, Sebastiaan Bol, Evelien M. Bunnik

Visualization: Rolando Garza, Evelien M. Bunnik

Writing – original draft: Rolando Garza, Evelien M. Bunnik

Writing – review & editing: Rolando Garza, Jeffrey Marchioni, Jared Honeycutt, Nicholas K. Hurlburt, Caroline Torres, Anakaren Garcia, Eva Loranc, Emily Yemington, Dalton Towers, Isaac Ssewanyana, Marie Pancera, Jason Lavinder, Prasanna Jagannathan, Bryan Greenhouse, Sebastiaan Bol, Evelien M. Bunnik

## CONFLICT OF INTEREST

The authors declare that they have no conflict of interest.

## DATA AVAILABILITY

All relevant data are within the paper and its supporting information files. Correspondence and requests for materials should be addressed to Evelien M. Bunnik.

